# Motor imagery and execution activate similar finger representations that are spatially consistent over time

**DOI:** 10.1101/2024.12.09.627466

**Authors:** Ingrid Angela Odermatt, Laura Schönberg, Caroline Heimhofer, Patrick Freund, Nicole Wenderoth, Sanne Kikkert

## Abstract

Finger representations in the sensorimotor cortex can be activated even in the absence of somatosensory input or motor output through mere top-down processes, such as motor imagery. While executed finger movements activate finger representations in the primary sensorimotor cortex that are spatially consistent over time within participants, the stability of top-down activated finger representations remains largely unexplored. Given the increasing use of top-down activated sensorimotor representations to both plan implantation of and control brain-computer interfaces, it is crucial to understand the stability of these representations. Here, we investigated the spatial consistency, and thereby reliability, of finger representations activated through motor imagery in the primary somatosensory and primary motor cortex over time. To assess this, participants performed imagined and executed individual finger movements in two 3T fMRI sessions that were ∼2 weeks apart. We observed highly consistent univariate finger-selective activity clusters and multivariate vertex-wise activity patterns within participants over time in both the motor imagery and motor execution task. Using a multivariate across-task decoding approach, we further found that motor execution and motor imagery activate similar finger representations in both the primary somatosensory and primary motor cortex. This demonstrates that motor imagery can be used to identify finger representations related to movement execution. Our findings not only validate the use of top-down processes for brain-computer interface planning and control, but also open up new opportunities for the development of sensorimotor training interventions that do not rely on overt movements.

## Introduction

The somatotopic organisation in the primary somatosensory cortex (S1) and primary motor cortex (M1) describes the correspondence between an area of the body and the specific area of the cortex representing that body part (Penfield & Boldrey, 1937). Neural finger representations have been of particular interest in this context given their fine-grained somatotopy (Besle et al., 2013; Ejaz et al., 2015; Kolasinski et al., 2016; Sanchez-Panchuelo et al., 2010; Sanders et al., 2023; Schellekens et al., 2018) and importance for fine motor skills (Xu et al., 2024). Their detailed somatotopic organisation has been extensively studied using functional magnetic resonance imaging (fMRI) (Janko et al., 2022). Traditionally, S1 and M1 finger representations are investigated using tactile stimulation or motor execution paradigms (Berlot et al., 2019; Sanders et al., 2023). Over the past years, evidence has accumulated demonstrating that neural representations of individual fingers in S1 and M1 can also be activated through top-down processes, i.e., without any overt motor output or somatosensory input. Specifically, finger representations have been detected in S1 by directing attention to individual fingers (Puckett et al., 2017), by anticipated tactile stimulation (Kassraian et al., 2023), by merely holding tactile information in working memory (Rabe et al., 2023), or by observed touch (Kuehn et al., 2018). Furthermore, finger representations are activated in both M1 and S1 during movement preparation (Ariani et al., 2022) or motor imagery (Odermatt et al., 2024). While it has been shown that finger representations activated through executed movements are spatially highly stable over time for up to six months (Ejaz et al., 2015; Kolasinski et al., 2016), the spatial consistency, and thus reliability, of top-down activated finger representations has not yet been explored.

The lack of a clear understanding regarding the reliability of top-down activated sensorimotor representations is surprising, given their growing neuroscientific and clinical importance. Indeed, the fact that top-down processes can activate representations of individual fingers has enabled researchers to study finger representations in populations in the absence of sensorimotor functions. Specifically, S1 and M1 finger representations have been detected through attempted finger movements of a missing hand after arm amputation or paralysed hand after cervical spinal cord injury (Kikkert et al., 2016, 2021; Wesselink et al., 2019). These preserved representations (Shah et al., 2023; van den Boom et al., 2021) are increasingly used to control brain-computer interfaces (BCIs) (Shah et al., 2024; Willsey et al., 2024). Using intracortical recordings in the sensorimotor cortex of cervical spinal cord injury patients with completely paralysed fingers, it has been demonstrated that it is possible to decode individual attempted and imagined finger movements even though actual motor output is completely lacking (Andersen & Aflalo, 2022; Guan et al., 2022, 2023; Jorge et al., 2020). Additionally, we have shown that it is possible to strengthen these top-down activated finger representations through motor imagery training in the absence of overt movements (Odermatt et al., 2024). Such top-down sensorimotor training interventions may allow to induce adaptive sensorimotor neuroplasticity in patients who are not (yet) able to perform overt movements (McFarland et al., 2020; Norman et al., 2018), for instance following a stroke or incomplete spinal cord injury.

To validate the use of and optimally leverage top-down activated sensorimotor representations for BCI control and training interventions, we investigated the reproducibility of fine-grained finger representations activated through motor imagery of three different fingers (thumb, index finger, and little finger) over time. To do so, we assessed finger representations in a motor imagery and a motor execution task in 16 healthy participants across two 3T fMRI sessions that were approximately two weeks apart. We investigated spatial consistency in S1 and M1 using two distinct approaches. First, we applied univariate analysis methods to quantify the spatial correspondence of finger-selective (or winner-take-all) maps. Second, we used multivariate analysis to examine the reproducibility of the full intricate activity patterns associated with individual (imagined) finger movements through a decoding approach. If finger representations activated through motor imagery (and motor execution) are spatially consistent over time, then the univariate analysis should reveal more spatial overlap of finger-selective activity across sessions for ‘same’ compared to ‘neighbouring’ or ‘non-neighbouring’ fingers. Furthermore, if motor imagery (and motor execution) elicit consistent activity patterns over time, the multivariate decoding approach should achieve high classification accuracy across sessions. In both the uni- and multivariate analysis, we expected higher consistency within participants than across participants. Given the intertwined nature of S1 and M1, we expected to obtain similar results in these areas. Additionally, due to the implicated role of the supplementary motor area (SMA) and the ventral (PMv) and dorsal premotor cortex (PMd) in top-down motor-related processes (Bruurmijn et al., 2017; Park et al., 2015; Pilgramm et al., 2016; Zabicki et al., 2017), we exploratively conducted uni- and multivariate analyses to characterise motor imagery finger representations in these secondary motor areas. Finally, we compared motor imagery finger representations to the representations activated through motor execution to test their neural similarity in M1 and S1.

## Results

### Univariate finger-selective maps activated through motor imagery are spatially consistent over time

Motor imagery of individual fingers revealed top-down activated finger-selective activity clusters in the primary sensorimotor cortex at the individual participant level based on winner-take-all finger maps that were minimally thresholded (Z > 2) (**Fig. 1**). These finger-selective activity clusters progressed from the thumb (laterally) to the little finger (medially), corresponding to the gradient of finger preference in characteristic hand maps (Besle et al., 2013; Kolasinski et al., 2016; Sanchez-Panchuelo et al., 2010; Sanders et al., 2023). Qualitative assessment of these motor imagery finger maps suggests some extent of reproducibility across sessions for most participants, but high variability across participants.

**Figure 1.**
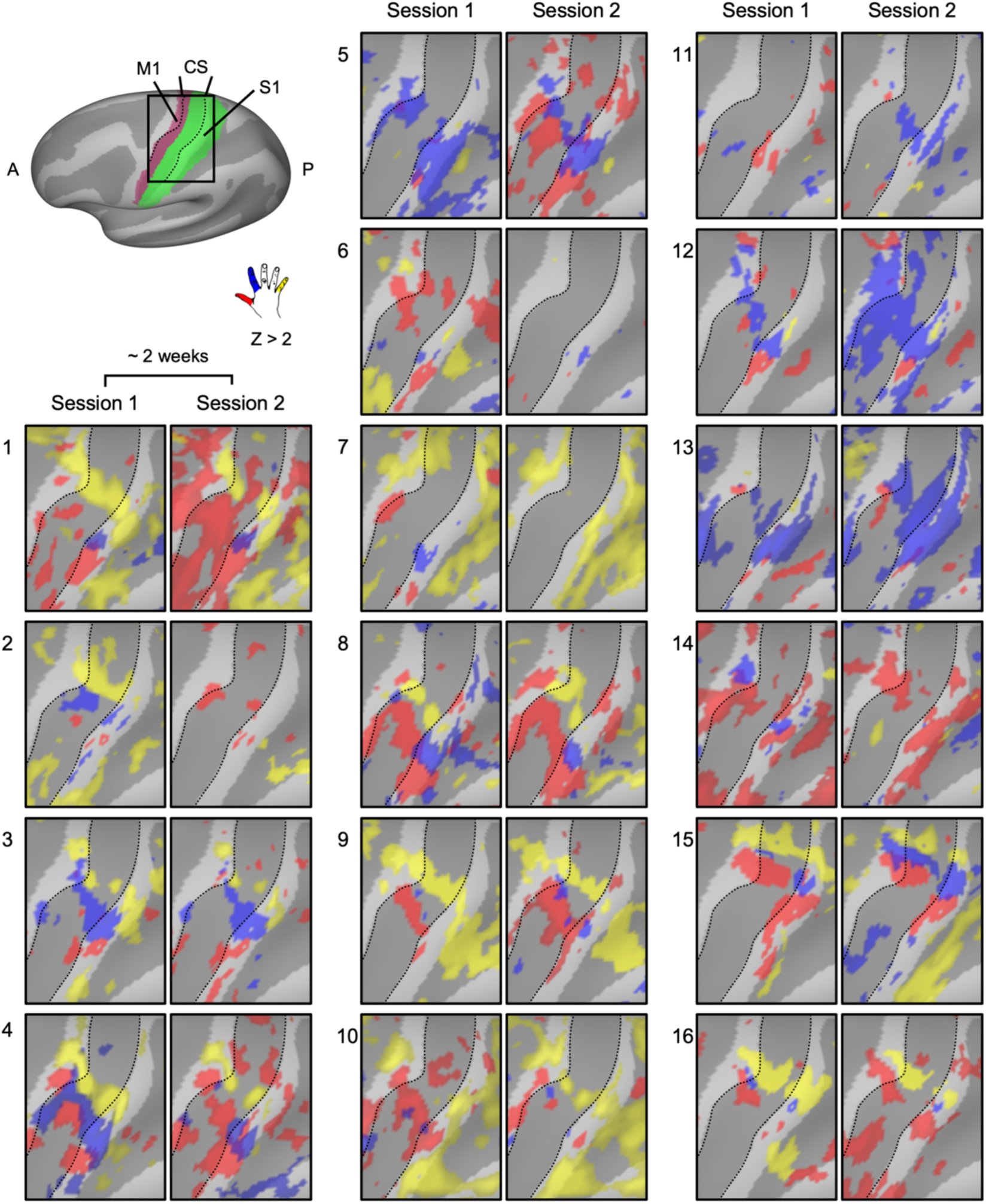
Minimally thresholded finger-selective motor imagery maps per participant across two fMRI sessions ∼2 weeks apart. Finger-selective motor imagery maps per participant (N = 16) visualised on a surface template brain with the sulcal pattern in dark grey. The top left shows the magnified area (black square), encompassing the hand area of the primary somatosensory (S1; green) and primary motor cortex (M1; purple). The colours of activity clusters refer to the winner-take-all finger contrasts thresholded at Z > 2 (red = thumb; blue = index finger; yellow = little finger). Typical finger-selectivity maps can be observed with a gradient of finger preference, progressing from the thumb (laterally) to the little finger (medially). While these characteristic aspects of finger maps are displayed for most participants, some finger representations are not visible in the minimally thresholded maps. A = anterior, P = posterior, CS = central sulcus.

To understand the stability, or spatial correspondence, of motor imagery finger maps, we then quantified the spatial overlap of the finger-selective (or winner-take-all) maps across sessions. Specifically, we calculated the Dice Overlap Coefficient (DOC) between every possible finger pair across both sessions within S1 and M1. In the main text, we focus on the results in S1 (**Fig. 2**), as we obtained similar results in M1 (**Supplementary Fig. 1)**. The explorative analysis for secondary motor areas is reported in **Supplementary Fig. 2**.

**Figure 2.**
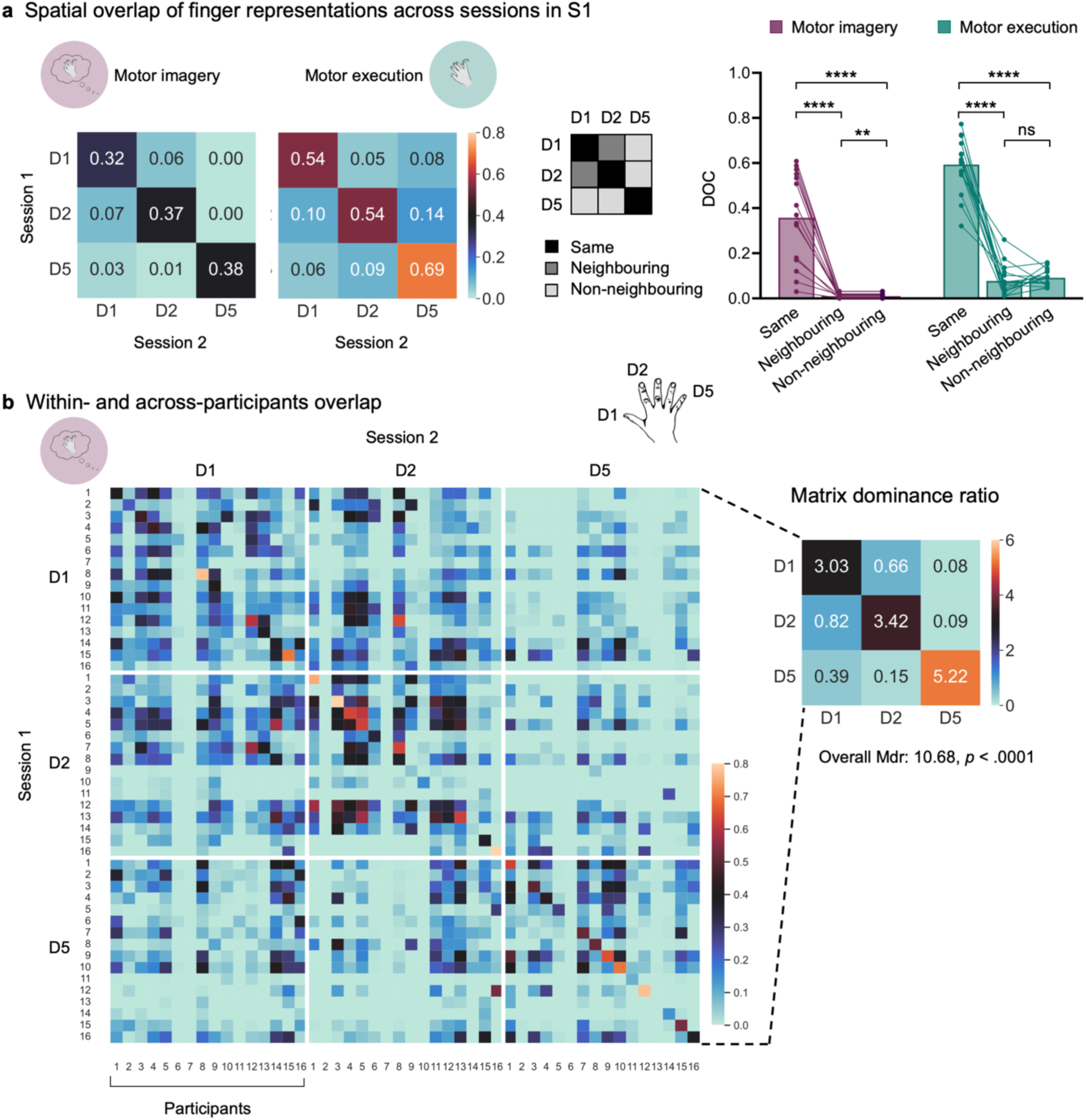
Finger-selective motor imagery maps in S1 are stable across sessions. **a**) Left: Spatial consistency, quantified using the Dice Overlap Coefficient (DOC), for all possible finger pairs across the two fMRI sessions. Results are averaged across all participants in the motor imagery and motor execution task based on the minimally thresholded (Z > 2) winner-take-all finger maps. Middle: The DOC were assigned to ‘same’ (diagonal of matrix: D1-D1, D2-D2, D5-D5), ‘neighbouring’ (D1-D2, D2-D1), or ‘non-neighbouring’ finger representations (D1-D5, D5-D1, D2-D5, D5-D2) for the statistical analysis depicted on the right. Dots depict data of individual participants. **** *p* < .0001; ** *p* < .01; ns = non-significant. **b**) Left: DOC comparing all combinations of individual finger representations across participants and sessions. The diagonal of the matrix represents the overlap of the same finger representations and participants across the two sessions. The 16 x 16 submatrices depicted by the white lines show the overlap for a specific finger pair for all participants. Right: Matrix dominance ratio (Mdr) for all submatrices based on the matrix on the left. The values > 1 on the diagonal reflect that the spatial overlap of the same finger representations was higher within-participant than across-participants. For neighbouring and non- neighbouring fingers there was no such higher spatial overlap within compared to across participants. From the Mdr matrix, an overall Mdr was calculated. A value > 1 indicates higher within-participant consistency than across-participants consistency. Bootstrapping revealed that the overall Mdr is significantly different from chance. D1 (digit 1) = thumb; D2 = index finger; D5 = little finger.

A DOC of 1 reflects a perfect overlap of two representations, while 0 indicates no overlap at all. In S1, our analysis revealed relatively high spatial consistency (DOC) for the ‘same’ motor imagery finger representations over time and low DOC for ‘neighbouring’ and ‘non-neighbouring’ finger representations (**Fig. 2a**), consistent with previously reported somatotopic finger maps (Kikkert et al., 2016, 2021; Kolasinski et al., 2016; Sanders et al., 2023). This was indicated in a significant *Task* by *Finger pair* interaction (χ^2^_(2)_ = 15.38, *p* < .001; main effect *Task*: χ^2^_(1)_ = 87.25, *p* < .0001; main effect *Finger pair*: χ^2^_(2)_ = 203.82, *p* < .0001). For the motor imagery task, the DOC for the same finger representations was higher compared to neighbouring (*t*_(85)_ = 7.09, *p* < .0001) and non-neighbouring finger representations (*t*_(85)_ = 9.12, *p* < .0001), and higher for neighbouring compared to non-neighbouring finger representations (*t*_(85)_ = 3.42, *p* = .003). This demonstrates a somatotopic gradient in the spatial correspondence across sessions and indicates that the finger maps activated through motor imagery were stable over time. Similarly, in the motor execution task, the DOC was higher for same compared to neighbouring (*t*_(85)_ = 10.73, *p* < .0001) and non-neighbouring finger representations (*t*_(85)_ = 11.20, *p* < .0001). However, there was no difference in the DOC for neighbouring and non-neighbouring finger representations (*t*_(85)_ = -0.72 *p* = .95), possibly due to the inclusion of only three fingers (i.e., thumb, index finger, little finger) in the analysis (see **Supplementary Fig. 3b** for DOC including all five fingers). When comparing the spatial consistency across the two tasks, we found lower DOC during motor imagery compared to motor execution for the same (*t*_(85)_ = -4.07, *p* = .003) and non-neighbouring (*t*_(85)_ = -9.27, *p* < .0001) finger representations, and no significant difference for neighbouring finger representations (*t*_(85)_ = -0.37, *p* = .95; for simplicity of visualisation not depicted in **Fig. 2a**).

Next, we compared the within-participant to across-participants consistency. To do so, we calculated the DOC across sessions for all possible finger and participant pairings. The across-participants DOC matrix (**Fig. 2b**) consists of submatrices representing each finger pair. If the spatial overlap of finger-selective activity clusters is higher within than across participants, then the values on the diagonal of the ‘same’ finger pair submatrices (i.e., D1-D1, D2-D2, D5-D5) should be higher than the values on the off-diagonal of these ‘same’ finger pair submatrices. As expected, within-participant DOC was higher than across-participants DOC for same fingers, as indicated by all matrix dominance ratios (Mdr) > 1 (Mdr D1 = 3.03, Mdr D2 = 3.42, Mdr D5 = 5.22; see diagonal of matrix on the right in **Fig. 2b**). By contrast, for both neighbouring and non-neighbouring finger pairs, all Mdrs were < 1 (see off-diagonal of matrix on the right in **Fig. 2b**). This indicates that for neighbouring and non-neighbouring finger-pairs, the spatial correspondence across sessions within a participant was not higher than the spatial correspondence across participants. Finally, the overall Mdr was calculated based on the Mdr scores of the submatrices in **Fig. 2b**. The resulting overall Mdr of 10.68 implies that spatial correspondence of the ‘same’ finger representations across sessions drives within-participant consistency. To investigate the likelihood of obtaining a Mdr score higher or equal to the obtained Mdr score merely by chance, we applied bootstrapping and found that the overall Mdr was significanty different from chance level (*p* < .0001), demonstrating higher within-participant than across-participants consistency of motor imagery finger representations.

Together, the DOC analyses suggest that, at the group level, the finger-selective activity clusters elicited through motor imagery can be studied using fMRI and follow a somatotopic organisation, similarly as finger maps activated through finger movement or tactile finger stimulation. Importantly, these somatotopic motor imagery finger maps are spatially consistent over weeks. However, at the individual participant level, the minimally thresholded finger maps did not reveal finger-selective activity clusters for each participant, session, and finger. If overall activity levels are (expected to be) low and the conditions are thought to be represented in distributed activity patterns, as suggested for individual (imagined) finger movements, then univariate analysis is suboptimal. Thus, we next used multivariate analysis to overcome these limitations.

### Imagined finger movements can be decoded with higher accuracy across sessions than across participants

Multivariate pattern analysis (MVPA) takes into account the full pattern of activity elicited by (imagined) finger movements. Here, we used a multivariate decoding approach to investigate whether imagined finger movements can be reliably predicted from the activity pattern in the S1 or M1 hand area across sessions, giving insight into the consistency of these activity patterns over time. To do so, we trained a linear support vector machine on the data of one session and tested it on the data of the other session. In the S1 hand area, we found that within-participant classifiers generalised well across sessions. Specifically, accuracy of this across-sessions decoding was significantly above the empirical chance level (*p* < .0001, **Fig. 3a**), and did not differ from within-session accuracy (*t*_(15)_ = 1.30, *p* = .21, BF_10_ = 0.52 indicating anecdotal evidence for the null hypothesis). This suggests that activity patterns associated with individual imagined finger movements are highly reproducible over fMRI sessions, demonstrating high within-participant consistency.

**Figure 3.**
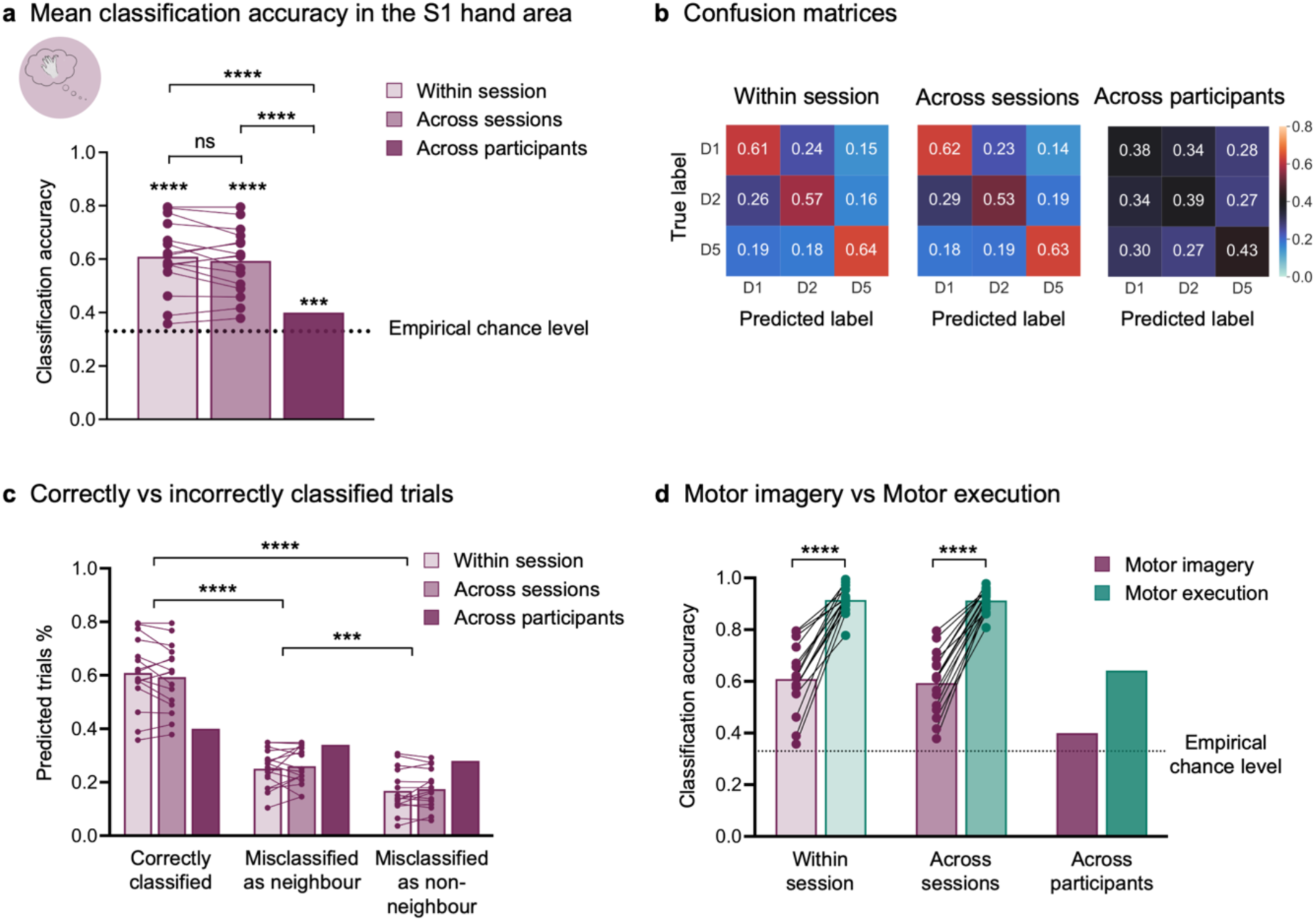
Decoding analysis of individual imagined finger movements reveals that finger-specific activity patterns are spatially consistent across sessions in the S1 hand area. **a**) Classification accuracy of individual fingers during motor imagery in the S1 hand area. Within-session classification accuracy depicts the average accuracy of a leave-one-run-out cross-validation performed separately for each participant and session. Across- sessions is based on the average accuracy score of a within-participant classifier trained on all trials of one session and tested on all trials of the other session. Across-participants classification accuracy shows the average accuracy of a leave-one-participant-out cross-validation using the data of all participants. Paired classical and Bayesian t-tests suggest that within-session and across-sessions accuracy do not differ significantly (BF_10_ = 0.52). One-sample t-tests demonstrate significantly higher classification accuracy for classifiers trained and tested within-participants compared to across-participants, indicating higher within-participant consistency of motor imagery finger represenations over time than across-participants. Asterisks on top of bars refer to the statistical difference of classification accuracy from the empirical chance level. **b**) Confusion matrices depicting the correctly classified trials (diagonal) and the misclassified trials (off-diagonal) in the S1 hand area. Each row refers to all trials of a finger, and the cells in a row show the % of predicted trials for each label. **c**) We assigned the scores of the confusion matrices in b) to ‘correctly classified’ (diagonal; true label – predicted label: D1-D1, D2-D2, D5-D5), misclassified as ‘neighbouring’ finger (D1-D2, D2-D1), and as ‘non-neighbouring’ (D1-D5, D5-D1, D2-D5, D5-D2) and found that significantly more trials were correctly classified than misclassified. From the misclassified trials, significantly more trials were wrongly predicted as a neighbouring than non-neighbouring finger. The across-participants classification scores are displayed merely for visualisation and were not included in the statistical analysis. The significance bars refer to post-hoc contrasts based on the main effect of *Prediction* (correctly classified, misclassified as neighbouring, misclassified as non-neighbouring finger), averaged over both levels of *Comparison* (within-session, across-sessions). **d**) Finger representations activated through motor imagery were less clearly distinguishable in the S1 hand area than when they were activated through motor execution. However, the generalisability of a classifier trained on trials of one session and tested on trials of the other session was comparable for motor imagery and motor execution. The significance bars refer to the main effect *Task* (motor imagery, motor execution). The across-participants scores are merely displayed for visualisation and were not included in the statistical analysis. The dotted lines represent the empirical chance level (i.e., 0.33) based on permutation testing. Dots depict data of individual participants. **** *p* < .0001; *** *p* < .001; ns = non-significant. D1 (digit 1) = thumb; D2 = index; D5 = little.

To investigate across-participants consistency in finger representations, we next tested the generalisability of a classifier to other participants and found that such an across-participants classifier was able to successfully decode individual fingers of participants whose data were unseen in the training (average accuracy in leave-one-participant-out cross-validation: 0.40, *p* < .001, empirical chance level: 0.33). However, classification accuracy was significantly lower compared to within-participant classification (within session vs. across participants *t*_(15)_ = 6.38, *p* < .0001; across sessions vs. across participants *t*_(15)_ = 6.37, *p* < .0001; **Fig. 3a**). These results demonstrate that within-participant consistency in activity patterns associated with individual finger motor imagery is significantly higher than across-participants consistency.

When comparing the true vs. the predicted labels in the decoding analyses (**Fig. 3b**), we found more misclassified trials for neighbouring fingers (i.e., thumb misclassified as index finger or vice versa), than for non-neighbouring fingers (i.e., thumb or index finger misclassified as little finger or vice versa; **Fig. 3c**; significant main effect *Prediction*: *F*_(2,90)_ = 187.22, *p* < .0001; non-significant main effect *Comparison*: *F*_(1,90)_ = 0.00, *p* = .99; non-significant interaction effect: *F*_(2,90)_ = 0.17, *p* = .85). Significantly more trials were correctly classified compared to misclassified as neighbouring (*t*_(75)_ = 14.68, *p* < .0001) or as non-neighbouring fingers (*t*_(75)_ = 18.26, *p* < .0001). From the wrongly classified trials, significantly more trials were misclassified as a neighbouring finger compared to another finger (*t*_(75)_ = 3.58, *p* = .0006), indicating higher similarity of vertex-wise activity patterns of neighbouring fingers compared to non-neighbouring fingers. Note that the values obtained from the across-participants classification are plotted for visualisation purposes only in **Fig. 3c** but were not considered in the statistical analysis as only a single value per *Prediction* level was available.

We then compared the classification accuracy scores obtained from decoding of fingers in the motor imagery vs. the motor execution task (**Fig. 3d**), and found, as expected, lower scores for imagined than executed finger movements (significant main effect *Task*: *F*_(1,45)_ = 364.59, *p* < .0001; non-significant main effect *Comparison*: *F*_(1,45)_ = 0.30, *p* = .59; non-significant interaction effect: *F*_(1,45)_ = 0.16, *p* = .69). While decoding of three fingers in the motor execution task lead to very high classification accuracies (mean within session: 0.92; mean across sessions: 0.91; empirical chance level: 0.33), including all five fingers lead to lower absolute values (mean within session: 0.63; mean across sessions: 0.58; empirical chance level: 0.20; **Supplementary Fig. 4**). These findings reveal that although motor imagery elicits less distinguishable activity patterns for individual fingers compared to motor execution, the reproducibility of these patterns across sessions does not differ significantly between the two tasks.

Similar results were obtained for the M1 hand area, as depicted in **Supplementary Fig. 5**. In secondary motor areas including SMA, PMd, and PMv we found that imagined and executed finger movements could be decoded significantly above chance, with most correctly classified trials for the thumb (**Supplementary Fig. 6**).

### Within-participant variability in vividness, motivation, and perceived effort do not predict across-sessions generalisability of a classifier that decodes imagined finger movements

In comparison to overt finger movements or tactile stimulation, motor imagery has a higher cognitive component that may increase variability in task performance both within and across participants. We thus investigated whether session-to-session variations in attentional and cognitive processes could explain the scores obtained from generalising a classifier across sessions. We calculated the absolute difference score from session 1 to session 2 for the subjective measures of vividness during motor imagery, perceived motivation, and effort. We then used these difference scores as fixed effects in a multiple linear regression to predict the average across-sessions motor imagery classification accuracy in the S1 hand area. This model did not reach significance (*F*_(3,12)_ = 1.87, *p* = .19, multiple *r*^2^ = .32; vividness: *t* = 0.46, *p* = .65; motivation: *t* = -1.21, *p* = .25, effort: *t* = -1.77, *p* = .10).

We then investigated whether the subjective measures would predict the within-session classification accuracy score, using the average of the subjective measures of both sessions as predictors, and found that motivation significantly predicted the average within-session classification accuracy (*t* = 2.44, *p* = .03). Vividness (*t* = -0.90, *p* = .39) and effort (*t* = .68, *p* = .51) did not reach significance (model: *F*_(3,12)_ = 5.05, *p* = .02, multiple *r*^2^ = .56).

### Neural similarity of finger representations activated through motor imagery vs. motor execution

Finally, we investigated the neural similarity between activity patterns elicited by individual imagined and executed finger movements. We used an across-task decoding analysis and trained a classifier on all data of one task in one session and tested it then on all data of the other task in either the same (within session) or other session (across sessions). This analysis revealed that we could successfully decode individual fingers across tasks, as indicated by classification accuracies significantly above the empirical chance level (all p < .0001; **Fig. 4**). We found that a classifier trained on motor imagery generalises better to motor execution, than vice versa (significant main effect of *Task*: *F*_(1,45)_ = 225.53, *p* < .0001). Within- and across-sessions classification accuracy did not differ (non-significant main effect *Comparison*: *F*_(1,45)_ = 0.09, *p* = .77; non-significant interaction effect: *F*_(1,45)_ = 0.07, *p* = .79). Similar results were obtained for the M1 hand area (**Supplementary Fig. 7**). These findings show that motor imagery and motor execution activate finger representations containing mutual information in their finger-specific vertex-wise activity patterns.

**Figure 4.**
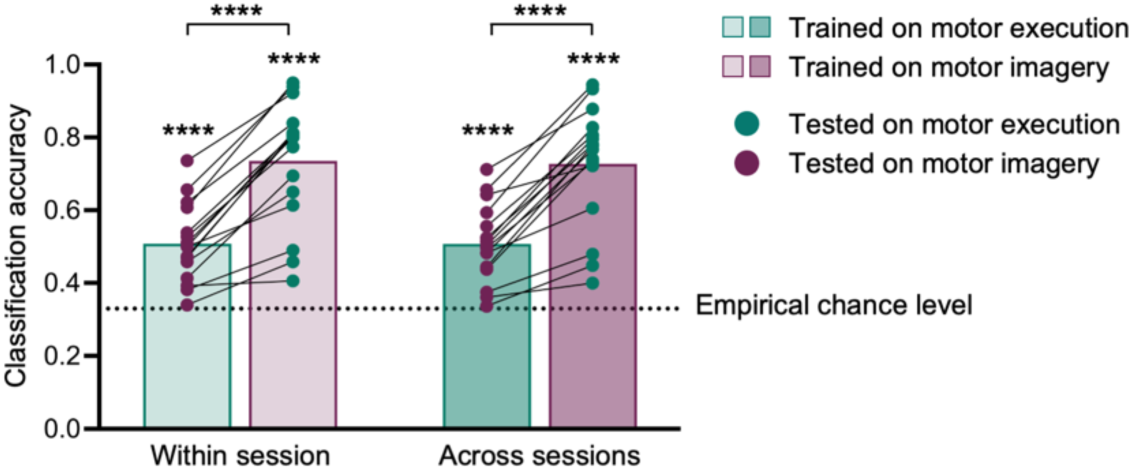
Across-task decoding analysis reveals neural similarity of finger representations activated through motor imagery and motor execution in the S1 hand area. Within-session classification accuracy depicts the average accuracy of a classifier trained on all data of one task (i.e., either motor imagery or motor execution) in one session and tested on all data of the other task in the same session, performed separately for each participant. Across-sessions shows the average accuracy of a classifier trained on all data of one task in one session and tested on all data of the other task in the other session. Asterisks on top of bars refer to the statistical difference of classification accuracy from the empirical chance level. The dotted lines represent the empirical chance level (i.e., 0.33) based on permutation testing. Dots depict data of individual participants. **** p < .0001; ns = non-significant.

## Discussion

In this study, we investigated the spatial consistency of finger representations activated through motor imagery over time. Our results show that both finger-selective activity clusters and the full intricate activity patterns associated with individual imagined finger movements were highly consistent across two fMRI sessions. The finding that motor imagery reliably activated finger representations is particularly striking given the highly unsupervised nature of the task. Notably, the level of consistency of finger representations activated through motor imagery was comparable to motor execution. Furthermore, an across-task decoding analysis revealed that motor imagery and motor execution activated finger representations that contained shared information in their underlying activity patterns, indicating neural similarity between these representations.

Using univariate and multivariate analyses, we found high within-participant consistency in finger representations activated through motor imagery in both S1 and M1. These findings align with the results of previous (Ejaz et al., 2015; Kolasinski et al., 2016) and the current study showing highly reproducible finger representations activated through motor execution, which we used as a benchmark for comparing our motor imagery findings. In line with the results from univariate analysis of motor execution finger representations reported in Kolasinski et al. (2016), we found that the spatial correspondence of finger-selective activity clusters in S1 for the same motor imagery finger representations across sessions was higher compared to different finger pairs, representing a hallmark of somatotopic organisation (Kikkert et al., 2016; Kolasinski et al., 2016; Sanders et al., 2023). While Kolasinski et al. (2016) employed a travelling-wave paradigm, we created finger maps based on a blocked paradigm. Travelling-wave paradigms are powerful in detecting finger maps based on gradients from the thumb to the little finger. In contrast, blocked paradigms allow to investigate more detailed somatotopic information that account better for the spatial overlap of finger representations (Besle et al., 2013; Kikkert et al., 2021). Nevertheless, even based on blocked paradigms, winner-take-all approaches have limitations as they strongly depend on the (arbitrarily) chosen threshold and the specific conditions (in our case, fingers) that were tested (Muret et al., 2022; Wesselink et al., 2019). Indeed, despite minimal thresholding, motor imagery finger maps did not show finger-selective activity clusters for all tested fingers in every participant. To circumvent arbitrarily thresholding of winner-take-all approaches, and to account for low activity levels during motor imagery and largely overlapping finger representations, we used a multivariate decoding approach and examined whether the finger with which motor imagery (or motor execution) was performed could be predicted from the full activity patterns of single trials. Our results confirm high reproducibility of these activity patterns associated with individual fingers across sessions for both motor imagery and motor execution.

For finger representations activated through motor imagery, we found lower absolute values in spatial consistency compared to motor execution. This was evident in lower spatial correspondence of the same finger representations across sessions in the univariate analysis and in lower classification accuracy when decoding fingers within- and across-session. Multiple factors may have contributed to this effect. First, it is likely that the activity levels evoked by motor execution were higher, resulting in a better signal-to-noise-ratio and thus more clearly distinguishable activity patterns associated with individual fingers. Second, the motor execution task was more standardised compared to the motor imagery task. For motor execution, participants performed paced button presses. If participants started a trial with a delay, we corrected the trial onsets and if they accidentally moved with the wrong finger, we assigned the trial to that finger. In contrast, the motor imagery task was unsupervised, self-paced, and without clear instruction towards the specific motor imagery strategies. Suggested strategies included pressing a button, playing the piano, typing on keyboard, moving the finger up and down or left and right, or imagine pulses in the finger muscle. We did not restrict participants to use only one motor imagery strategy and left it open to change strategies from trial to trial or session to session (see **Supplementary Table 1** for instructions and self-reported strategies). These task differences may have resulted in higher variability in the motor imagery performance and thus resulting activity patterns. Third, top-down processes may be more affected by task compliance and attentional and cognitive processes compared to motor execution tasks, additionally contributing to more variability in motor imagery finger representations. Indeed, attention can greatly modulate finger representations measured with fMRI (Puckett et al., 2017) and vividness of imagined action has been shown to be reflected in more distinguishable top-down activated representations of hand actions (Zabicki et al., 2019). We found that session-to-session variability of perceived vividness of motor imagery, motivation and how exhausting the motor imagery task was perceived, did not predict the generalisability of a classifier across sessions, suggesting that top-down activated finger representations may be robust against small session-to-session fluctuations in attentional and cognitive processes. However, higher self-reported motivation scores were associated with higher decoding performance within session, indicating that motivation (and thus, task compliance) may indeed play a role. In summary, it is likely that an interplay of all processes contributed to lower classification accuracy and lower spatial overlap of finger-selective activity for motor imagery compared to motor execution. Nevertheless, higher classification accuracies may be achievable through training, as we have demonstrated that neurofeedback training for mental finger individuation results in more distinguishable activity patterns of individual imagined finger movements (Odermatt et al., 2024).

In the across-task decoding analysis, we found neural similarity between activity patterns elicited by imagined and executed finger movements. We have previously reported that motor imagery activates similar finger representations as motor execution in the primary sensorimotor cortex (Odermatt et al., 2024). Here, we extend this finding by showing high spatial consistency over time across tasks. Notably, the across-task decoding analysis performed better when trained on motor imagery and tested on motor execution, than vice versa. We speculate that the lower signal-to-noise-ratio and higher within-session variability in activity patterns elicited by motor imagery compared to motor execution (as described in the previous paragraph), may have contributed to this finding. If motor execution of individual fingers elicits highly consistent activity patterns across trials, a decoder trained on this data may exhibit decreased performance when tested on the more variable activity patterns of motor imagery (Hebart & Baker, 2018). Sensorimotor representations activated through motor imagery are thought to be more sparse and widespread than motor execution patterns (Zabicki et al., 2017). While motor execution patterns may be encompassed within the broader motor imagery patterns, and thus be successfully predicted by a decoder trained on motor imagery patterns, the reverse case may be more challenging. These findings of our across-task decoding analysis add to the knowledge on neural equivalence of imagined and executed movements, which is relevant for the development of motor imagery training interventions aiming to improve sensorimotor functions.

In both motor imagery and execution, we found similar patterns of spatial consistency in S1 and M1, reaffirming their tight inter-connections and co-involvement not only in executed (Ogawa et al., 2019; Sanders et al., 2023; Umeda et al., 2019; Wesselink et al., 2019) but also imagined finger movements. While S1 activity during motor execution is modulated by the sensory feedback from finger movements, it has been shown that top-down processes elicit activity in the form of predicted sensory consequences prior to (and in the complete absence of) sensory input (Ariani et al., 2022; Bashford et al., 2021; Gale et al., 2021; Jafari et al., 2020; London & Miller, 2013; Umeda et al., 2019). This top-down induced activity, or corollary discharge, in S1 is highly somatotopic with a topographic arrangement of individual fingers, while neuronal populations in M1 may be less selective towards individual fingers (Arbuckle et al., 2022; Ejaz et al., 2015; Graziano et al., 2002; Hluštík et al., 2001; Kakei et al., 1999; Schellekens et al., 2018; Schieber, 2001). Studying M1 and S1 separately is interesting due to their differences in somatotopic organisation. However, the exact delineation of M1 and S1 is challenging with 3T fMRI given the spatial resolution. Most previous studies investigated uni- and multivariate characteristics of finger representations using 7T fMRI (Besle et al., 2013, 2014; Gooijers et al., 2022; Kolasinski et al., 2016; Sanchez-Panchuelo et al., 2010; Sanders et al., 2023; Wesselink et al., 2022) achieving a spatial resolution of ∼ 1mm^3^ (as opposed to 2.2 mm^3^ in the present study). However, our results relating to the motor execution task are in line with studies using ultra-high field fMRI (Ejaz et al., 2015; Kolasinski et al., 2016), indicating that the similar findings obtained in S1 and M1 are not merely effects of the lower spatial resolution in our study, but likely attributable to the co-involvement of S1 and M1 during (imagined) finger movements.

Our results have implications for technologies relying on top-down processes to activate sensorimotor representations such as the control of BCIs, including neuroprosthetic devices. While top-down processes during fMRI assessments have been used for presurgical planning to optimally locate BCI target regions (Downey et al., 2024; Leinders et al., 2023), the reliability of these approaches to activate finger representations has not yet been explicitly studied. For BCI localisation, attempted movements are typically employed, as they were shown to result in higher blood oxygen level dependent (BOLD) activity compared to motor imagery (Hotz-Boendermaker et al., 2008) and better discrimination of specific representations using intracortical recordings (Vargas-Irwin et al., 2018).

However, it is likely that some patients are unable to perform attempted movements. Our findings suggest that in these cases motor imagery can be used instead to reliably reveal finger maps. Additionally, our results validate the use of 3T fMRI to study and visualise top-down finger representations, which makes is more accessible in research and clinical settings than its 7T equivalent. The stability of top-down sensorimotor representations across sessions is promising for the use of decoding approaches that are intended for longer-term use. Finally, despite rather high variability across participants, similar as for finger representations activated through motor execution, we observed that motor imagery finger representations share some information across participants. This was evidenced in a decoding approach that successfully generalised across participants and implies that technologies focusing on decoding individual fingers, such as BCIs for fine-motor control, may benefit from pre- training with large datasets of multiple participants to improve performance.

Together, our results highlight the use of fMRI and multivariate analysis that takes into account the full activity pattern as a reliable method to study (top-down activated) finger representations in sensorimotor areas, also if a clearly ordered somatotopic organisation is lacking and activity levels are relatively low. We show that motor imagery, a mere top-down process, activates consistent finger representations over time. Additionally, we demonstrate that finger representations activated through motor imagery and motor execution contain mutual information. These findings not only validate the use of top-down processes for brain-computer interface control, but also have implications for the development of top-down training interventions targeting sensorimotor functions without relying on overt movements.

## Methods

The data used in this manuscript was published previously in Odermatt et al. (2024). The focus of this already published manuscript lay on changes in finger representations of a neurofeedback intervention group compared to a control group that merely served to control for test-retest effects. In the current manuscript we explore the data of the control group in greater detail. The motor imagery and execution tasks, as well as fMRI acquisition and preprocessing in the current manuscript are therefore identical to Odermatt et al. (2024). The relevant sections are reiterated below for the reader’s convenience.

### Participants

16 volunteers (age (mean ± SD): 26.4 ± 2.7 years; 8 females) participated in the current study. Inclusion criteria were: No use of medication acting on the central nervous system, no neurological and psychiatric disorders, right-handed according to the Edinburgh Handedness Inventory (Oldfield, 1971), normal or corrected-to-normal vision, and no MRI contraindications. All research procedures were approved by the Cantonal Ethics Committee Zurich (BASEC Nr. 2018-01078) and were conducted in accordance with the declaration of Helsinki. Written informed consent was provided by all participants prior to study onset.

### Experimental sessions

In this study, we report data collected in two identical fMRI sessions that took place on the same time of the day and at an interval of approximately two weeks (mean ± SD: 13.8 ± 6.2 days; range: 7 – 32 days). Prior to the first fMRI session, participants were screened for their ability to perform kinaesthetic motor imagery using the kinaesthetic subscale of the Movement Imagery Questionnaire – Revised second version (MIQ-RS) (Gregg et al., 2010; Thomschewski et al., 2017). None of the recruited participants exhibited a score below the defined cut-off that was set to 1 SD below the mean reported in Gregg et al. (2010). In an experimental session prior to the first fMRI session, participants already performed kinaesthetic motor imagery of individual fingers (right thumb, index, or little finger) while we measured surface electromyography (EMG) in the left and right thumb, index, and little finger and applied transcranial magnetic stimulation (TMS) single- and paired-pulse testing protocols. In this pre-fMRI session, participants trained to imagine movements without making any actual movements or muscle contractions by receiving online visual feedback indicating if the EMG activity in any of the finger muscles exceeded 10 µV (Odermatt et al. 2024). Importantly, there was no experimental session between the two fMRI sessions.

### fMRI tasks

In the two fMRI sessions, we assessed brain activity during imagined and executed right hand finger movements. Participants viewed a fixation cross centred on a screen through a mirror mounted to the head coil. To restrict head motion, we used padded cushions. We first assessed four motor imagery runs using a blocked design with the conditions ‘Thumb’, ‘Index’, ‘Little’ and ‘Rest’ (block length = 7.5 s; each condition presented 12 x per run in counterbalanced order; 9 min 8s per run; **Fig. 5**). The verbal task instructions replaced the fixation cross for the duration of the trial. After each trial, the fixation cross reappeared for a jittered period of 3-4 s during which participants were instructed to rest, i.e., as during the rest trials. This additional rest period was included to facilitate switching between different fingers during the motor imagery task. We instructed participants to imagine, as vividly as possible, how it would feel to move the cued finger (see **Supplementary Table 1a** for verbatim instructions). Additionally, we reminded them to not contract any finger muscles, as trained in the pre-fMRI session with EMG activity control. An experimenter visually controlled for finger movements inside the scanner room. If any movements were detected, we stopped the run, instructed the participant to refrain from executing finger movements, and repeated the run. At the end of each motor imagery run, participants used a button box to move a cursor on a visual analogue scale (ranging 0-100) to rate, per finger, how vividly they felt they imagined the movements (verbal anchors: *not vivid at all*; *felt as vivid as during real movements*). At the end of the last motor imagery run, participants additionally rated their motivation (*not motivated at all*; *perfectly motivated*), focus on motor imagery (*not focused at all*; *perfectly focused*), and effort (*not exhausting at all*; *extremely exhausting*) during the session.

**Figure 5.**
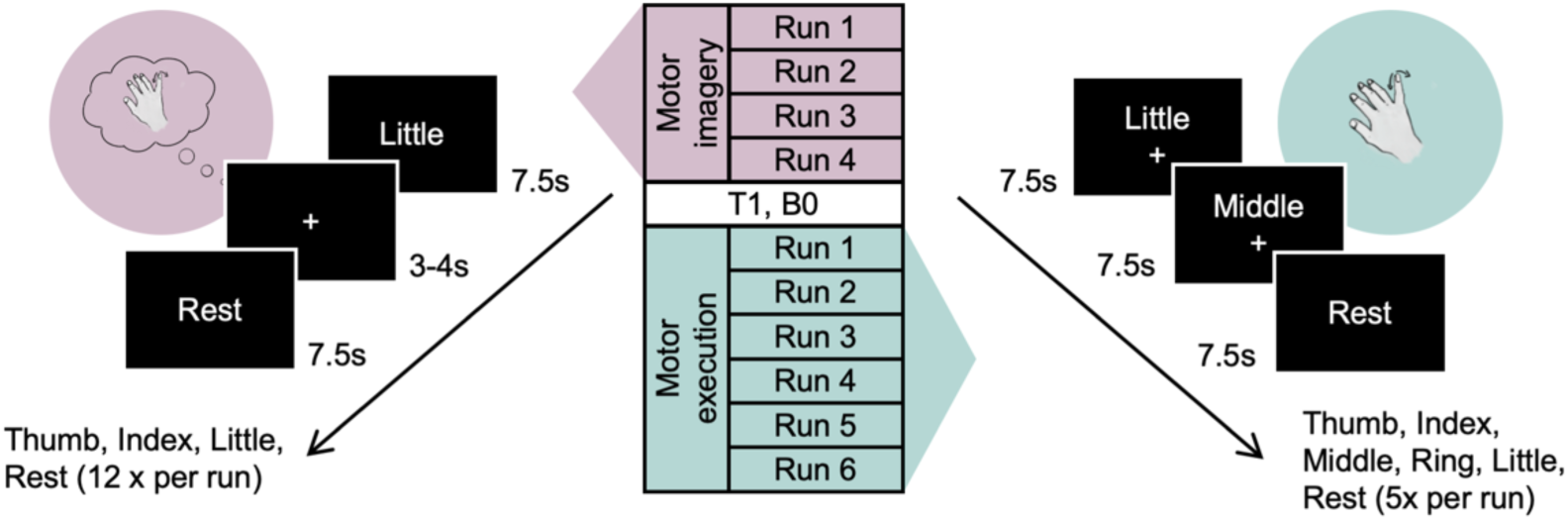
Experimental setup of one fMRI session. In each fMRI session, four motor imagery runs (purple) were acquired using a blocked paradigm. In each block, participants were visually cued to imagine individual finger movements by the words ‘Thumb’, ‘Index’, ‘Little’, or ‘Rest’, presented in a counterbalanced order. After each trial, an additional rest period (3-4s) to facilitate switching between different fingers was indicated by a fixation cross. After acquiring a T1 and B0 sequence, six motor execution runs were assessed, using an identical blocked paradigm as for motor imagery but without additional rest periods between the movement trials. The participants’ right index, ring, middle and little fingers were placed on the buttons of a four-button response box, with the thumb placed on the side of the box. Participants were visually cued by the words ‘Thumb’, ‘Index’, ‘Middle’, ‘Ring’, ‘Little’, or ‘Rest’ displayed above the fixation cross to perform paced button presses with the corresponding finger (or to tap the side of the button box with the thumb) or to rest. The pace was instructed by the fixation cross blinking at 1.4 Hz. In the rest trials, no fixation cross was displayed.

Following the motor imagery runs, we assessed six motor execution runs that consisted of a paced button press task and included the conditions ‘Thumb’, ‘Index’, ‘Middle’, ‘Ring’, ‘Little’, and ‘Rest’. We used a similar blocked paradigm as for motor imagery but did not provide any additional jittered rest periods between the movement trials (block length = 7.5 s; each condition presented 5 x per run in counterbalanced order; 4 min 5 s per run; **Fig. 5**). The verbal task instructions cueing the condition appeared above the fixation cross. Participants placed their right index, ring, middle and little fingers on the buttons of a four-button response box, with the thumb placed on the side of the box. We instructed participants to press the button with the cued finger (or tap on the side of the button response box with the thumb) every time the fixation cross blinked (1.4 Hz).

### fMRI data acquisition

We used a 3T Siemens Magnetom Prisma scanner with a 64-channel head-neck coil (Siemens Healthcare, Erlangen, Germany) to acquire fMRI data. For the anatomical T1-weighted images, we used a Magnetization Prepared Rapid Gradient Echo (MPRAGE) protocol with the following acquisition parameters: 160 sagittal slices, resolution = 1.1 x 1.1 x 1 mm^3^, field of view (FOV) = 240 x 240 x 160 mm^3^, repetition time (TR) = 2300 ms, echo time (TE) = 2.25 ms, flip angle = 8°. For the task-fMRI data acquisition we used a multiband echo-planar-imaging (EPI) sequence covering the whole brain and the cerebellum with the following acquisition parameters: 66 transversal slices, resolution = 2.2 mm^3^ isotropic, FOV = 210 x 210 x 145 mm^3^, TR = 846 ms, TE = 30 ms, flip angle = 56°, acceleration factor = 6, and echo spacing = 0.6 ms. We acquired 636 and 278 volumes for each of the motor imagery and motor execution runs, respectively. To measure B0 deviations we used a fieldmap with the same resolution and slice angle as the EPI sequence and the following acquisition parameters: TR = 649 ms, TE1 = 4.92 ms, TE2 = 7.38 ms.

### Data preprocessing and co-registration

For preprocessing and co-registration of fMRI data we used tools from FSL v5.0.7 (http://fsl.fmrib.ox.ac.uk/fsl) and applied the following preprocessing steps to the fMRI data using FSL’s Expert Analysis Tool (FEAT): motion correction using MCFLIRT (Jenkinson et al., 2002), brain extraction using the automated brain extraction tool (BET) (Smith, 2002), spatial smoothing using a 3 mm full-width at half-maximum (FWHM) Gaussian kernel, and high-pass temporal filtering with a 100 s cut-off. Using BET and/or Advanced Normalization Tools (ANTs) v2.3.5 (http://stnava.github.io/ANTs) we removed non-brain tissue from the T1-weighted images of two fMRI sessions and created binarized masks of the extracted brains. We then performed image co-registration in separate, visually inspected steps. For each participant, we created a mid-space, i.e., an average space, between the T1-weighted images of both sessions and its binarized brain masks. We then used the mid- space brain mask to brain extract the mid-space T1-weighted image. By using this T1-weighted mid- space for co-registration we ensured that the extent of reorientation required in the registration from functional to structural data was equal in both fMRI sessions. We then aligned functional data to the brain-extracted T1-weighted mid-space, initially using six degrees of freedom and the mutual information cost function, and then optimised using boundary-based registration (BBR) (Greve & Fischl, 2009). To correct for B0 distortions, we constructed a fieldmap for B0 unwarping and added it to the registration. For one participant that we took out of the scanner for a brief break during the first fMRI session, we applied the fieldmap only to the functional runs that were acquired with the same head position as the fieldmap.

### General Linear Models

#### Univariate analysis

To assess univariate task-related activity of motor imagery and execution, we carried out time-series statistical analysis per run using FMRIB’s Improved Linear Model (FILM) with local autocorrelation, as implemented in FSL’s FEAT. We defined one regressor of interest for each individual finger (i.e., for motor imagery: thumb, index, and little finger; for motor execution: thumb, index, middle, ring, and little finger) and obtained parameter estimates using a GLM based on the gamma hemodynamic response function (HRF) and its temporal derivatives. For motor execution, we examined the recorded button presses of each trial to control whether participants used the instructed finger to tap the button and adjusted the finger movement regressors if necessary: If the switch to the next cued finger in a new trial was delayed, then we adjusted the corresponding block lengths and the onset of the next trial. If the button of a non-instructed finger was pressed during a trial, then we adjusted the regressors such that the trial was assigned to this non-instructed, moving, finger. To reduce noise, we added nuisance regressors for the six motion parameters (rotation and translation along the x, y, and z-axis), and white matter and cerebrospinal fluid time series to the GLM. We further assessed the data for excessive head motion and scrubbed volumes with an estimated absolute displacement greater than 1.1 mm (i.e., half of the functional voxel size; maximum percentage of volumes scrubbed in a run = 2.16 %).

We defined contrasts for each finger > other fingers and averaged across runs at the individual participant level using fixed-effect analysis to obtain the winner-take-all z-statistic maps, i.e. finger-selective activity clusters, for each finger in native space. For motor execution, we created contrasts for all five fingers vs. the other fingers, and additionally, to compare the univariate analysis with motor imagery, for thumb > index and little finger, index > thumb and little finger, and little finger > thumb and index finger.

### Multivariate analysis

For multivariate pattern analysis (MVPA), we computed single-trial parameter estimates using an HRF-based first-level GLM in SPM12 (http://www.fil.ion.ucl.ac.uk/spm/) using SPM’s default parameters, and disabled implicit masking based on the global mean intensity. The design matrix consisted of individual regressors for each motor imagery and motor execution trial based on the run-wise regressors described in the previous paragraph. This resulted in 48 parameter estimates per finger, session, and participant for motor imagery, and 30 for motor execution (unless the motor execution trials were adjusted as described in the previous paragraph). Additionally, we included the temporal derivatives and the same nuisance regressors as in the GLM for univariate analysis.

To directly compare the MVPA results in motor imagery with motor execution, we computed a second GLM for the motor execution runs with individual regressors for all thumb, index, and little finger trials, and regressors of no interest that included all middle and ring finger trials.

### Surface projections

For optimal co-registration and functional signal alignment across participants, we projected all contrast and parameter estimate maps to the cortical surface, using FreeSurfer v6.0 (Dale et al., 1999; Fischl, 2012) and Connectome Workbench v1.3.2 (Marcus et al., 2011). First, we reconstructed the cortical surface of each individual participant’s T1-weighted mid-space image. Following visual inspection of the pial and white matter reconstruction, we corrected any pial surface errors by manually adjusting the individual participant’s brain mask image. Then, we projected the winner-take-all z-statistic maps obtained from each individual participant’s fixed-effect analysis and the single-trial parameter estimate maps to the individual surface using cortical-ribbon mapping. Finally, we resampled the maps to the HCP fsLR-32k surface template brain (Van Essen et al., 2012).

### Regions of interest

Regions of interest (ROIs) were anatomically defined on the fsLR-32k surface template brain. For the univariate analysis (spatial overlap analysis), we defined contralateral M1 as Brodmann area (BA) 4, and S1 as BA 1, 2, 3a and 3b from the parcellation of Glasser et al. (2016). For MVPA, we used the contralateral M1 and S1 hand area masks covering 2 cm above and below the anatomical hand knob area (as made available by the Diedrichsen Lab on https://github.com/DiedrichsenLab/fs_LR_32). Our exploratory analyses included secondary motor areas, namely supplementary motor area (SMA), ventral premotor cortex (PMv), and dorsal premotor cortex (PMd). As these secondary brain regions were slightly overlapping with the M1 mask obtained from the Glasser atlas, we removed any overlapping voxels from all ROIs to avoid a voxel being assigned to multiple ROIs.

### Spatial overlap analysis

The Dice Overlap Coefficient (DOC) (Dice, 1945) was used to quantify the spatial consistency of univariate finger-selective (or winner-take-all) motor imagery and motor execution maps across sessions and participants. The DOC calculates the spatial overlap between two representations relative to the total area of these representations and varies from 0 (no spatial correspondence between finger representations) to 1 (perfect spatial correspondence between finger representations). Where A and B are the areas of two finger representations, the DOC is expressed as:

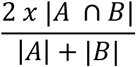

We closely followed previously described procedures (Gozzi et al., 2024; Kikkert et al., 2016; Kolasinski et al., 2016; Sanders et al., 2023). We minimally thresholded (Z > 2) the winner-take-all maps to obtain finger-selective activity clusters and masked these clusters using the ROIs (S1, M1, SMA, PMd, or PMv). We then used the in-vertex volume between the white matter and pial surfaces to calculate the volume covered by the thresholded representations and the resulting DOCs.

### Within participant

To assess within-participant consistency of univariate finger maps, we calculated the DOC for all possible finger pairings across the two fMRI sessions per participant. We then categorised the finger pairs in ‘same’, ‘neighbouring’ and ‘non-neighbouring’. A benchmark for finger- selectivity identified in univariate finger maps is greater DOC for the same finger representations than for neighbouring finger pairs, followed by non-neighbouring finger pairs (Kikkert et al., 2016; Sanders et al., 2023). Here, we investigated whether there is such spatial correspondence in the finger-selectivity across two fMRI sessions for motor imagery finger representations and whether it differed from motor execution. If spatial correspondence is high across two sessions, the DOC is higher for the same finger representations across sessions compared to neighbouring and non-neighbouring representations. To investigate how the choice of threshold influences the spatial overlap results, the DOC was further calculated as a function of Z-thresholds ranging from 1.4 to 3.2 (see **Supplementary Fig. 2 and Supplementary Fig. 3**).

### Within vs. across participants

To investigate the spatial correspondence of finger maps across participants, we calculated the DOC for all possible finger pairs and participants across sessions and constructed an across-participants DOC comparison matrix (48 x 48 for motor imagery and motor execution with three fingers; 80 x 80 for motor execution with five fingers), similar to Kolasinski et al., (2016). The across-participants matrix consisted of submatrices (16 x 16) containing the DOC for the 16 participants for each possible finger pair. We then calculated the matrix dominance ratio (Mdr) for each submatrix (i.e., each finger pair) by dividing the average of the diagonal (i.e., DOCs of the same participant) to the average of the off-diagonal (i.e., DOCs across different participant pairs). Values > 1 indicate higher overlap of finger representations within the same participants than across participants. We then computed the overall Mdr by calculating a higher-order Mdr of the Mdr scores in the submatrices. Here, a value > 1 indicates a matrix dominance for spatial correspondence of ‘same’ finger representations compared to ‘different’ (i.e., neighbouring and non-neighbouring) finger pairs. To obtain a p-value indicating significance of the overall Mdr value, we applied bootstrap resampling (n = 10’000) to the across-participants DOC comparison matrix.

### Multivariate pattern analysis

While the (univariate) spatial overlap analysis gives insight into the consistency of winner-take-all finger-specific activity clusters across sessions and participants, it does not account for the full activity pattern associated with individual (imagined) finger movements. In contrast, MVPA allows to examine the spatial consistency of the full intricate activity patterns associated with individual (imagined) finger movements. Here, we used multivariate classification analysis to decode the instructed finger during motor imagery (or motor execution) from the single-trial parameter estimates in a specific ROI (S1 hand area, M1 hand area, SMA, PMd, or PMv), using the scikit-learn python library (Abraham et al., 2014). To prepare the data set, we extracted the single-trial parameter estimates corresponding to the vertices in a ROI using nibabel (Brett et al., 2024). We then scaled the data across all motor imagery (or motor execution) runs for each session and participant separately with the StandardScaler from scikit-learn to remove the mean and scale to unit variance within each vertex. Next, we performed the classification analysis using a Support Vector Machine (SVM) with a linear kernel and default parameters of C = 1 and l2 regularisation.

### Within session

Within each session, and separately for each participant, we conducted a leave-one-run-out cross-validation. We evaluated the classifier performance based on the classification accuracy averaged across cross-validation folds and labels. To compare the actual and predicted labels, we averaged the confusion matrices obtained from each cross-validation fold. To define chance level, we generated a null distribution based on 1000 random permutations of the trial labels. We then computed an empirical p-value by dividing the number of permutation-based classification accuracies that were greater than or equal to the true score +1, by the number of permutations + 1 as implemented in scikit-learn. Finally, we averaged the classification accuracies, confusion matrices and empirical p-values across the two sessions for each participant.

### Across sessions

To investigate whether a classifier trained on one session within a participant generalised to data acquired in another session, we performed an across-sessions decoding analysis. For that, we fitted a linear SVM on all data of the first session and tested it on all trials of the second session, and vice versa. To determine the empirical chance level and p-values, we shuffled the labels of the test set using 1000 permutations. We then averaged the two classification accuracies, confusion matrices and p-values for each participant.

### Across participants

To test whether a classifier may generalise across participants, we conducted a leave-one-participant-out cross-validation pooling the data of both sessions together. This analysis resulted in only one value indicating classification accuracy on ‘group level’.

### Across tasks

Finally, to examine the neural similarity of activity patterns associated with individual fingers elicited by the motor imagery vs. the motor execution task, we trained a classifier on data of one task and tested it on data of the other task. We performed this across-tasks decoding analysis within- and across-sessions.

### Data analysis

For statistical analyses we used R v.4.3.1 (R Core Team, Vienna, Austria) and JASP v. 0.18.3 (JASP Team 2024, Netherlands). To determine statistical significance of classification accuracy at group level, i.e., whether classification accuracy is significantly above the empirical chance level, we combined the empirical p-values across participants using Fisher’s method (Fisher, 1992). To compare within-session to across-session accuracy, we used paired t-tests (or non-parametric Wilcoxon signed-rank tests if normality was violated according to the Shapiro-Wilk test). To compare within-session or across-session accuracy to across-participants accuracy (that consisted of only one value per ROI), we used one-sample t-tests.

To examine the effects and interaction of tasks and conditions on DOC and classification accuracy, we used lme4 (Bates et al., 2015) for linear mixed-effects models. We defined all factors (*Task* (motor imagery, motor execution); *Finger-pair* (same, neighbour, non-neighbour); *Comparison* (within-session, across-sessions); *Prediction* (correctly classified, misclassified as neighbouring finger, misclassified as non-neighbouring finger) as fixed effects and participant as random effect. For each model, we evaluated the expected against observed residuals for uniform distribution using the R package DHARMA (Hartig, 2022). If violations were detected (which was the case for the models on DOC in both S1 and M1), we used glmmTMB (Brooks et al., 2017) instead, which allows the dispersion parameter of the linear mixed-effects model to vary with the different levels of the factors. If the linear mixed-effects models revealed significant main effects or interaction, we computed post-hoc contrasts using emmeans (Lenth, 2024).

To examine whether subjective measures of attentional and cognitive states (i.e., vividness of motor imagery averaged across the three fingers and the four runs, motivation, focus and effort; see **Supplementary Table 2** for differences in the subjective measures across the two sessions) were associated with within-session classification accuracy, we performed linear regressions on the average within-session classification accuracy using the average of the subjective measures reported in both sessions. To assess potential multicollinearity of fixed effects, we examined the VIF (Variance Inflation Factor). Due to high multicollinearity, we excluded the predictor with the highest VIF (focus: VIF = 9.76), and rerun the model, resulting in low (i.e., < 5; James et al., 2021) multicollinearity of predictors (vividness = 2.87, effort = 2.87, motivation = 2.59). To evaluate whether changes in the subjective measures predict the generalisibility of a classifier across sessions we calculated session 2 minus session 1 difference scores per subjective measure and included these as fixed effects in the model. The average score of the across-sessions classification accuracy was the dependent variable. Multicollinearity was again high (focus: VIF = 25.96). After excluding focus, VIFs were < 5 for the remaining predictors (vividness = 1.87, effort = 1.11, motivation = 1.76).

The significance level alpha was set to 0.05 for all inferential tests. We corrected the p-values for multiple comparisons using the Bonferroni Holmes method within each ROI. For non-significant results, we further used Bayesian tests with default settings in JASP to provide evidence for or against the null hypothesis and reported the Bayes factor BF_10_ following conventional cut-offs (Dienes, 2014): BF_10_ < 0.1: strong evidence for the null hypothesis; BF_10_ < .33: moderate evidence for the null hypothesis, BF_10_ < 1: anecdotal evidence for the null hypothesis, BF_10_ = 1: no evidence.

## Acknowledgements

We thank all participants of the study, the Swiss Center for Musculoskeletal Imaging (SCMI) at Balgrist Campus for support with fMRI data acquisition, and Lukas Graz for his advice on statistical analysis. This project is supported by the Swiss National Science Foundation Grant 32003B_207719, and the National Research Foundation, Prime Minister’s Office, Singapore under its Campus for Research Excellence and Technological Enterprise (CREATE) program (FHT), S.K. is supported by the Swiss National Science Foundation Ambizione Grant (PZ00P3_208996).

## Supplementary Material

**Supplementary Figure 1.**
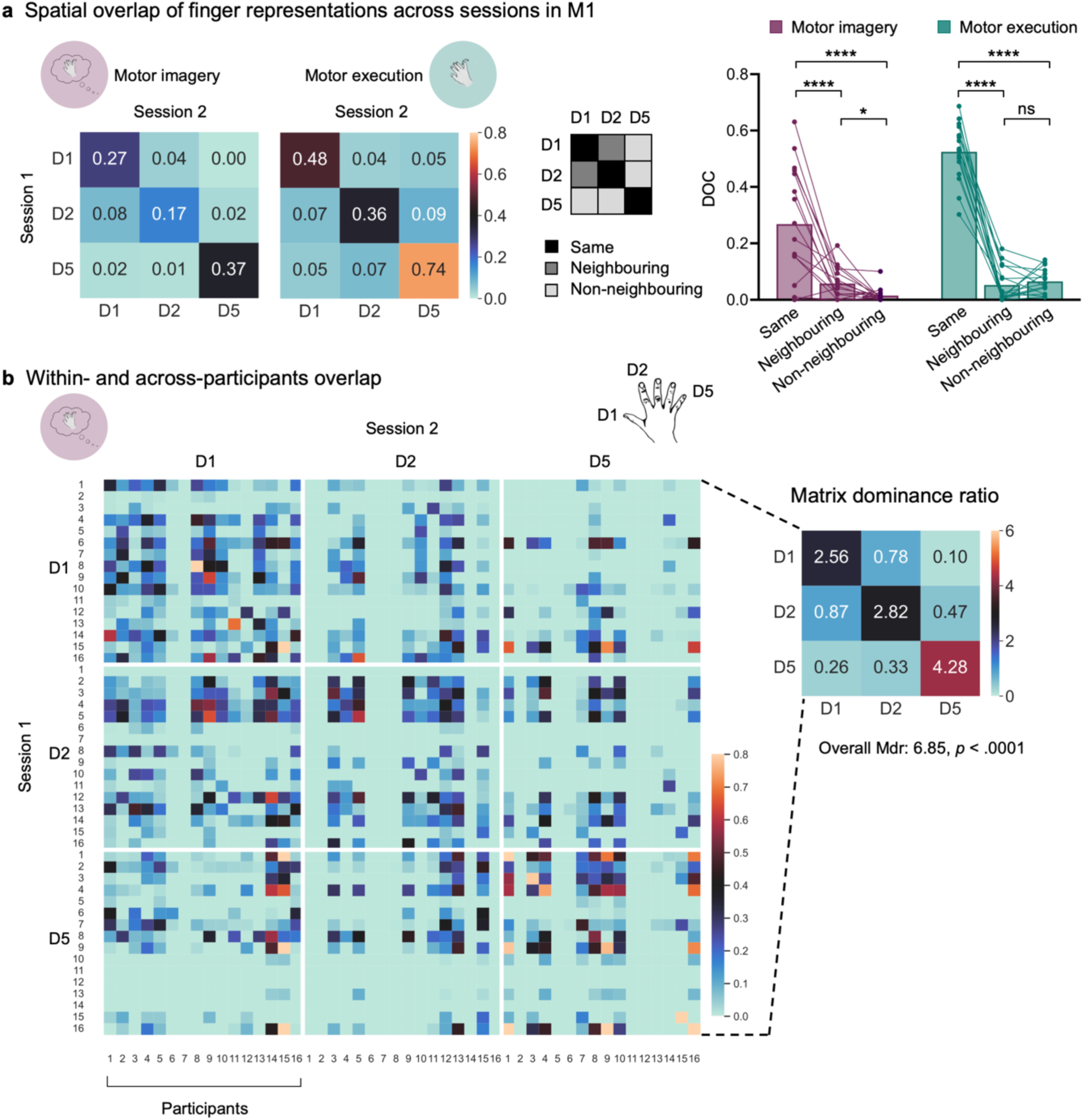
Finger-selective motor imagery maps in M1 are stable across sessions. **a**) Left: Spatial consistency, quantified using the Dice Overlap Coefficient (DOC), for all possible finger pairs across the two fMRI sessions. Results are averaged across all participants in the motor imagery and motor execution task based on the minimally thresholded (Z > 2) winner-take-all finger maps. Middle: The DOC were assigned to ‘same’ (diagonal of matrix: D1-D1, D2-D2, D5-D5), ‘neighbouring’ (D1-D2, D2-D1), or ‘non-neighbouring’ finger representations (D1-D5, D5-D1, D2-D5, D5-D2) for the statistical analysis depicted on the right. Our analysis revealed a significant 2 (*Task*: motor imagery, motor execution) by 3 (*Finger pair*: same, neighbouring, non-neighbouring) interaction (χ^2^ _(2)_ = 23.94, p < .0001; main effect *Task*: χ^2^ _(1)_ = 20.72, *p* < .0001; main effect *Finger pair*: χ^2^ _(2)_ = 186.21, *p* < .0001). For motor imagery we found higher DOC for same vs. neighbouring (*t*_(85)_ = 5.11, *p* < .0001) and non-neighbouring (t_(85)_ = 6.36, *p* < .0001), and higher DOC for neighbouring vs. non-neighbouring (*t*_(85)_ = 2.75, *p* = .02) finger representations. For motor execution we found higher DOC for same vs. neighbouring (*t*_(85)_ = 12.59, *p* < .0001) and non-neighbouring (*t*_(85)_ = 12.66, *p* < .0001), but not for neighbouring vs. non-neighbouring (*t*_(85)_ = -0.90 *p* = .74) finger representations. Comparing motor imagery with execution, we found that the DOC was lower for the same (*t*_(85)_ = -4.85, *p* = .003) and non-neighbouring (*t*_(85)_ = -4.59, *p* < .0001) finger representations during motor imagery but not for neighbouring finger representations (*t*_(85)_ = 0.28, *p* = .78; for simplicity of visualisation not depicted in the figure. Dots depict data of individual participants. **** *p* < .0001; * *p* < .05; ns = non-significant. **b**) Left: DOC comparing all combinations of individual finger representations across participants and sessions. The diagonal of the matrix represents the overlap of the same finger representations and participants across the two sessions. The 16 x 16 submatrices depicted by the white lines show the overlap for a specific finger pair for all participants. Right: Matrix dominance ratio (Mdr) for all submatrices based on the matrix on the left. The values > 1 on the diagonal reflect that the spatial overlap of the same finger representations was higher within-participants than across-participants. For neighbouring and non-neighbouring fingers there was no such higher spatial overlap within compared to across participants. From the Mdr matrix, an overall Mdr was calculated. A value > 1 indicates higher within-participant consistency than across-participants consistency. Bootstrapping revealed that the overall Mdr is significantly different from chance. D1 (digit 1) = thumb; D2 = index finger; D5 = little finger.

**Supplementary Figure 2.**
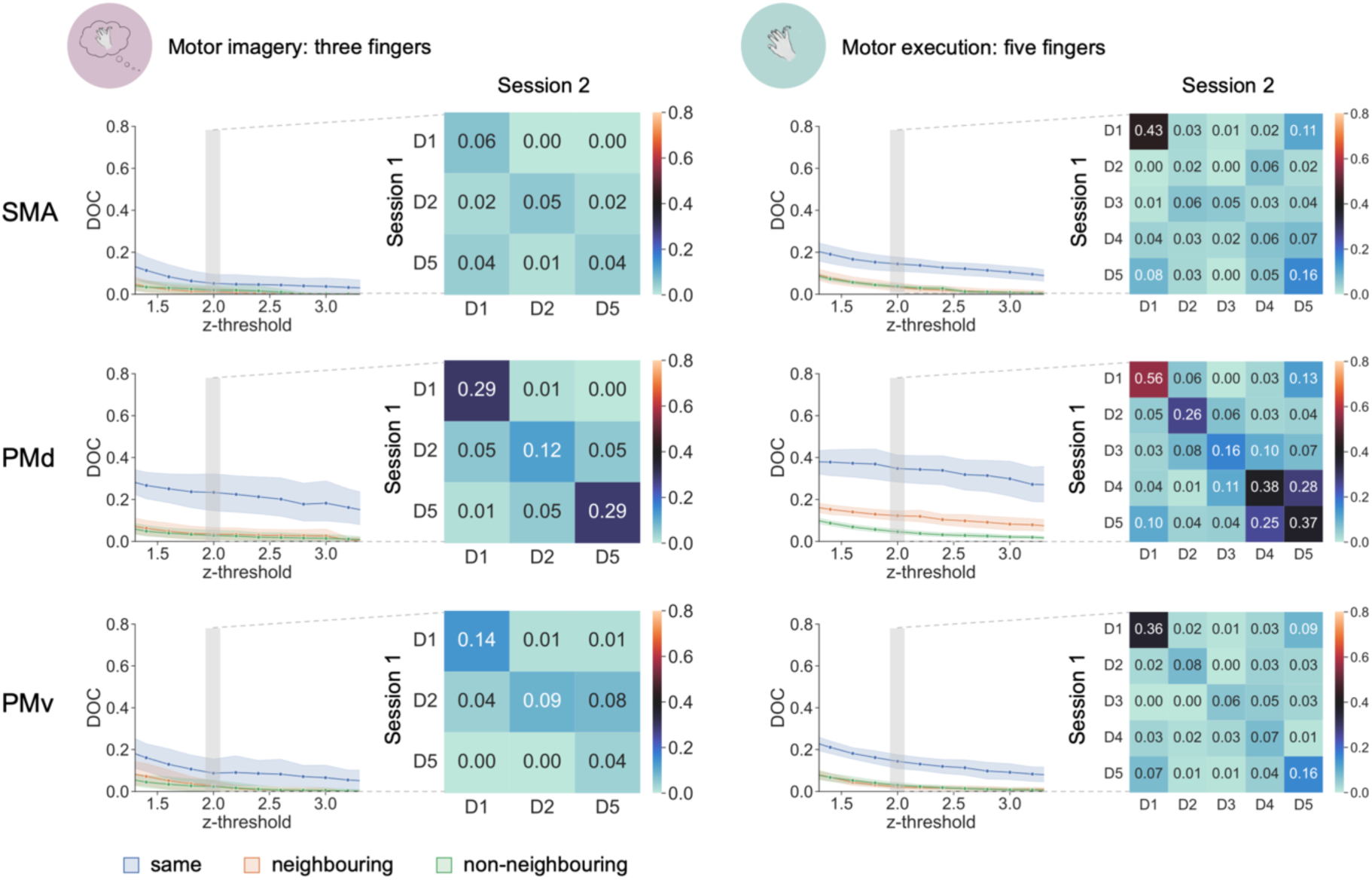
Dice Overlap Coefficient (DOC) across two fMRI sessions in secondary motor areas. DOC at different Z-thresholds assigned to ‘same’ (D1-D1, D2-D2, D5-D5; blue), ‘neighbouring’ (D1-D2, D2-D1; orange), or ‘non-neighbouring’ (D1-D5, D5-D1, D2-D5, D5-D2; green) for motor imagery (left) and motor execution (right). Dots depict the group average, shaded areas the 95% confidence interval. The matrices represent the DOC for all possible finger pairs across the two fMRI sessions, averaged over all participants based on the minimally thresholded (Z > 2) winner-take-all finger maps. D1 (digit 1) = thumb; D2 = index finger; D3 = middle finger; D4 = ring finger; D5 = little finger; SMA = supplementary motor area; PMd = dorsal premotor cortex; PMv = ventral premotor cortex.

**Supplementary Figure 3.**
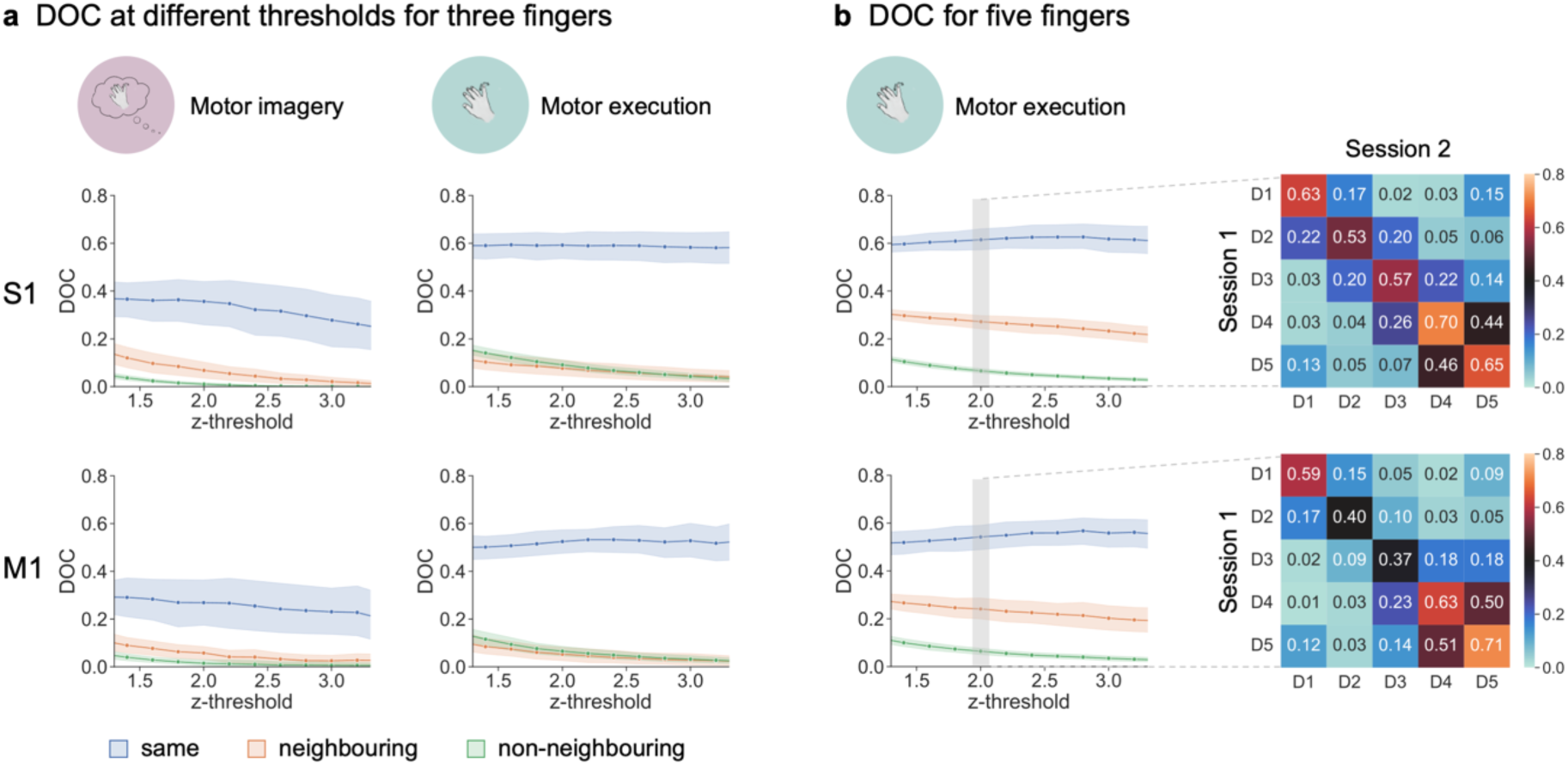
Dice Overlap Coefficient (DOC) across two fMRI sessions for different thresholds in M1 and S1. **a**) DOC at different Z-thresholds for three finger representations, assigned to ‘same’ (D1-D1, D2-D2, D5-D5; blue), ‘neighbouring’ (D1-D2, D2-D1; orange), or ‘non-neighbouring’ (D1-D5, D5-D1, D2-D5, D5-D2; green), separately for motor imagery and motor execution. Dots depict the group average, shaded areas the 95% confidence interval. The DOC matrices for all possible finger pairs are reported elsewhere (S1: Fig. 2a; M1: **Supplementary Fig. 1a**) **b**) Left: DOC at different Z-thresholds including all five fingers for motor execution. Right: DOC for all possible finger pairs across the two fMRI sessions, averaged over all participants in the motor execution task based on the minimally thresholded (Z > 2) winner-take-all finger maps. D1 (digit1) = thumb; D2 = index finger; D3 = middle finger; D4 = ring finger; D5 = little finger.

**Supplementary Figure 4.**
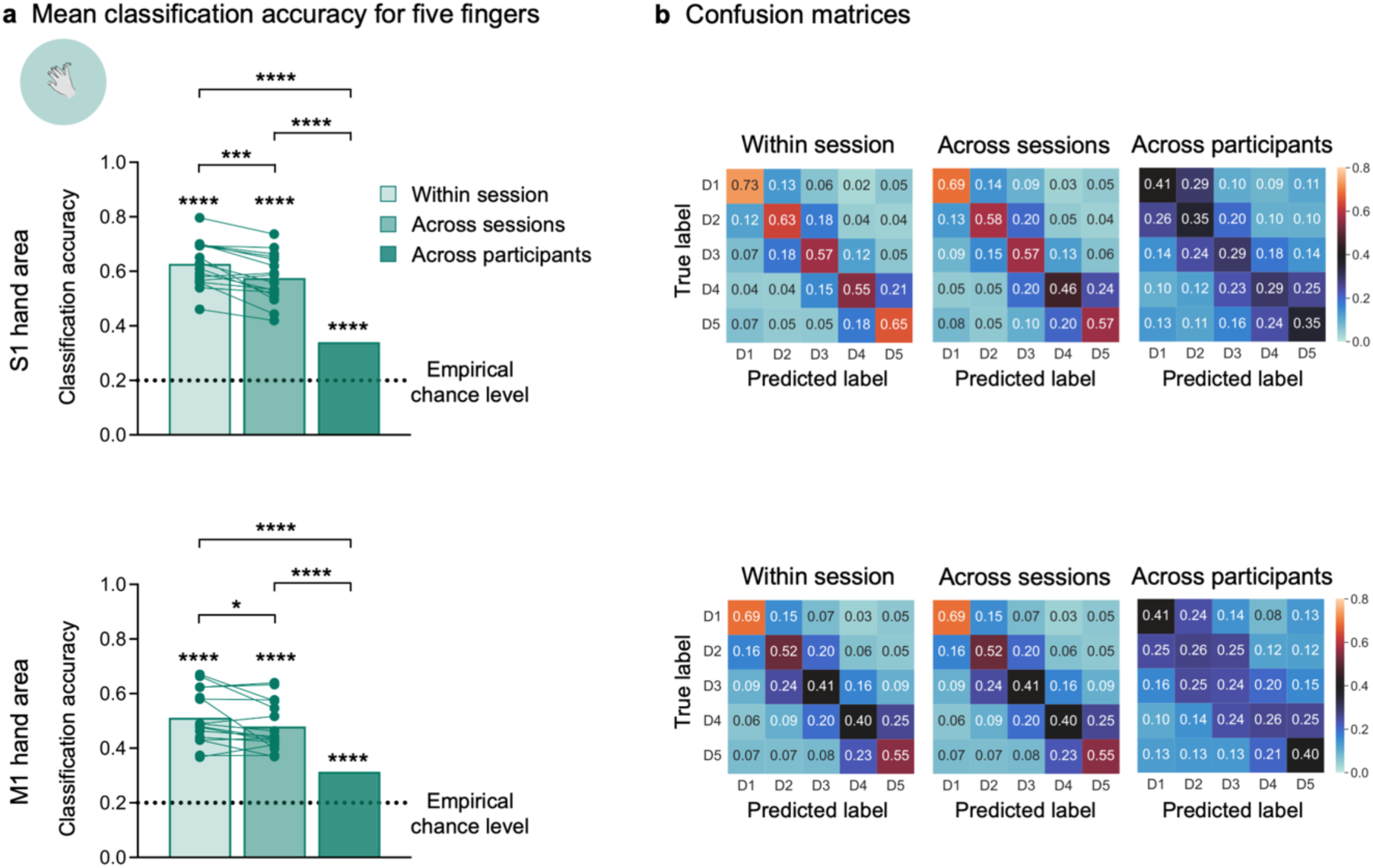
Decoding analysis of individual fingers in a motor execution task in the S1 and M1 hand area. **a**) Classification accuracy of five individual fingers during motor execution in the S1 hand area (top) and M1 hand area (bottom). Within-session classification accuracy depicts the average accuracy of a leave-one-run-out cross-validation performed separately for each participant and session. Across-sessions is based on the average accuracy score of a within-participant classifier trained on all trials of one session and tested on all trials of the other session. Across-participants classification accuracy shows the average accuracy of a leave-one-participant-out cross-validation using the data of all participants. Classification accuracy is significantly higher within-session than across-sessions based on paired t-tests (S1 hand area: *t*_(15)_ = 4.49, *p* = .0004; M1 hand area: *t*_(15)_ = 2.15, *p* = .048). One-sample t-tests demonstrate significantly higher classification accuracy for within-participant classifiers compared to across-participants decoding for both the S1 hand area (within-session vs. across-participants: *t*_(15)_ = 13.69, *p* < .0001; across-sessions vs. across-participants: *t*_(15)_ = 10.61, *p* < .0001) and M1 hand area (within-session vs. across-participants: *t*_(15)_ = 7.98, *p* < .0001; across-sessions vs. across-participants: *t*_(15)_ = 7.43, *p* < .0001). Asterisks on top of bars refer to the statistical difference of classification accuracy from the empirical chance level. **b**) Confusion matrices depicting the correctly classified trials (diagonal) and the misclassified trials (off-diagonal) in the S1 hand area (top) and M1 hand area (bottom). Each row refers to all trials of a finger, and the cells in a row show the % of predicted trials for each label. **** *p* < .0001; *** *p* < .001; * *p* < .05. D1 (digit1) = thumb; D2 = index finger; D3 = middle finger; D4 = ring finger; D5 = little finger.

**Supplementary Figure 5.**
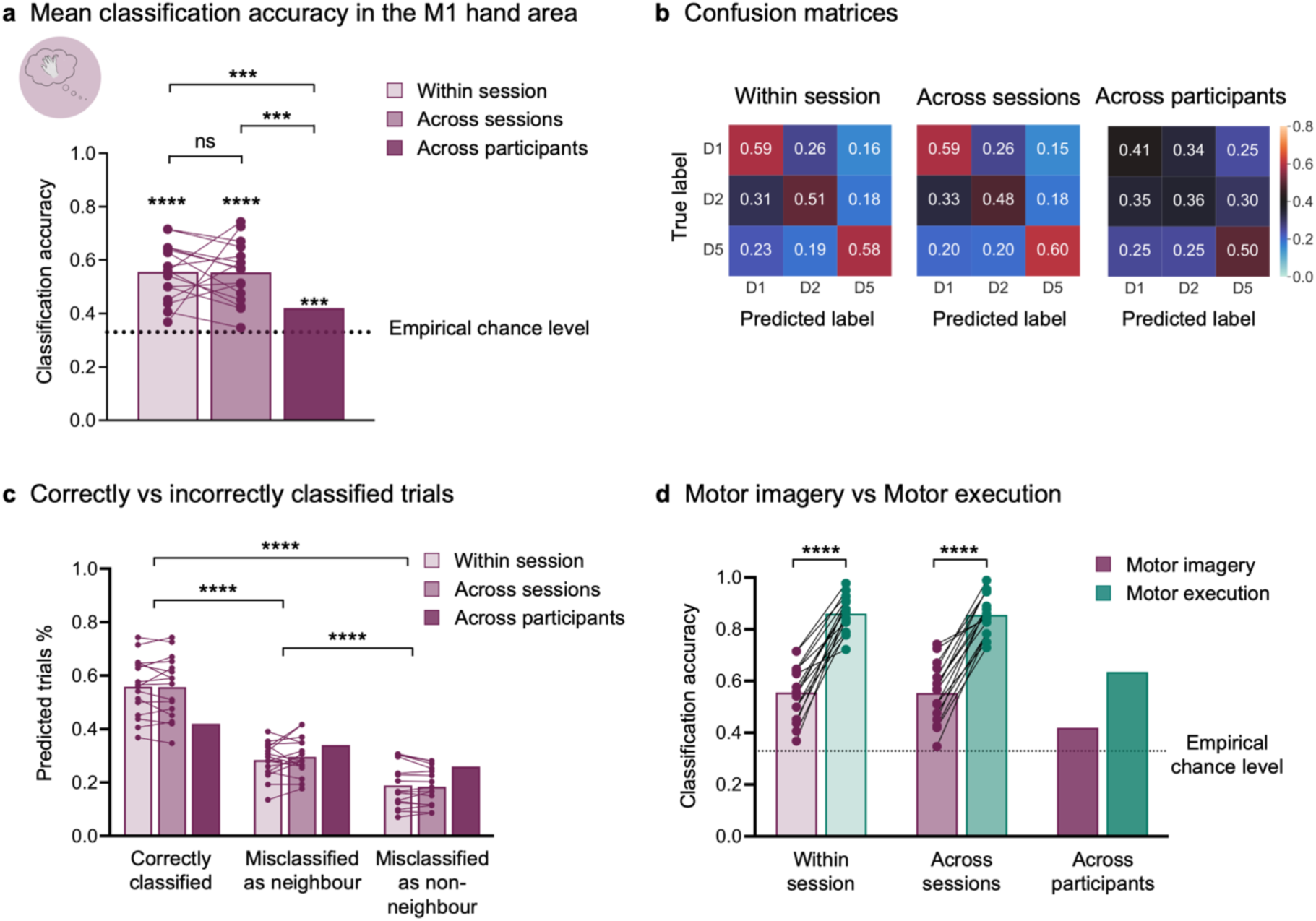
Decoding analysis of individual imagined finger movements reveals that finger-specific activity patterns are spatially consistent across sessions in the M1 hand area. **a**) Classification accuracy of individual fingers during motor imagery in the M1 hand area. Within-session classification accuracy depicts the average accuracy of a leave-one-run-out cross-validation performed separately for each participant and session. Across-session is based on the average accuracy of a within-participant classifier trained on all trials of one session and tested on the other session. Across-participant classification accuracy shows the average accuracy of a leave-one-participant-out cross-validation using the data of all participants. Paired classical and Bayesian t-tests suggest that within-session and across-sessions accuracy do not differ (*t*_(15)_ = 0.15, *p* = .88; BF_10_ = 0.26 indicating moderate evidence for the null hypothesis). One-sample t-tests demonstrate significantly higher classification accuracy for classifiers trained and tested within-participants compared to across-participants (within-session vs. across-participants: *t*_(15)_ = 4.99, *p* = .0005; across-sessions vs. across-participants: *t*_(15)_ = 4.68, *p* = .0006), indicating higher within-participant consistency of motor imagery finger represenations over time than across-participants. Asterisks on top of bars refer to the statistical difference of classification accuracy from the empirical chance level. **b**) Confusion matrices depicting the correctly classified trials (diagonal) and the misclassified trials (off-diagonal) in the S1 hand area. Each row refers to all trials of a finger, and the cells in a row show the % of predicted trials for each label. **c**) By assigning the scores of the confusion matrices to ‘correctly classified’ (diagonal; true label – predicted label: D1-D1, D2-D2, D5-D5), misclassified as ‘neighbouring’ finger (D1-D2, D2-D1), and as ‘non-neighbouring’ (D1-D5, D5-D1, D2-D5, D5-D2), we found that significantly more trials were correctly classified than misclassified (correctly classified vs. misclassified as neighbouring: *t*_(75)_ = 12.26, *p* < .0001; correctly classified vs. misclassified as non-neighbouring: *t*_(75)_ = 16.99, *p* < .0001). From the misclassified trials, significantly more trials were wrongly predicted as a neighbouring finger than non-neighbouring (*t*_(75)_ = 4.73, *p* < .0001). The scores for across-participants are displayed for visualisation merely and were not included in the statistical analysis. The significance bars refer to post-hoc contrasts based on the main effect of *Prediction* (*F*_(2,90)_ = 153.84, *p* < .0001), averaged over both levels of *Comparison* (within-session, across-sessions) following a linear-mixed effects model (main effect *Comparision*: *F*_(1,90)_ = 0.01, *p* = .92; *Prediction* by *Comparison* interaction: *F*_(2,90)_ = 0.08, *p* = .92). **d**) Finger representations activated through motor imagery were less clearly distinguishable in the M1 hand area than when they were activated through motor execution. However, the generalisability of a classifier trained on trials of one session and tested on trials of the other session was comparable for motor imagery and motor execution(linear mixed-effects model with fixed effects *Task* (motor imagery, motor execution) and *Comparison* (within-session, across-sessions); main effect *Task*: *F*_(1,45)_ = 497.10, *p* < .0001; main effect *Session*: *F*_(1,45)_ = 0.07, *p* = .79; *Task* by *Comparison* interaction: *F*_(1,45)_ = 0.03, *p* = .87). The significance bars refer to the main effect *Task*. The across-participants scores are merely displayed for visualisation and were not included in the statistical analysis. The dotted lines represent the empirical chance level (i.e., 0.33) based on permutation testing. Dots depict data of individual participants. **** *p* < .0001; *** *p* < .001; ns = non-significant. D1 (digit 1) = thumb; D2 = index; D5 = little.

**Supplementary Figure 6.**
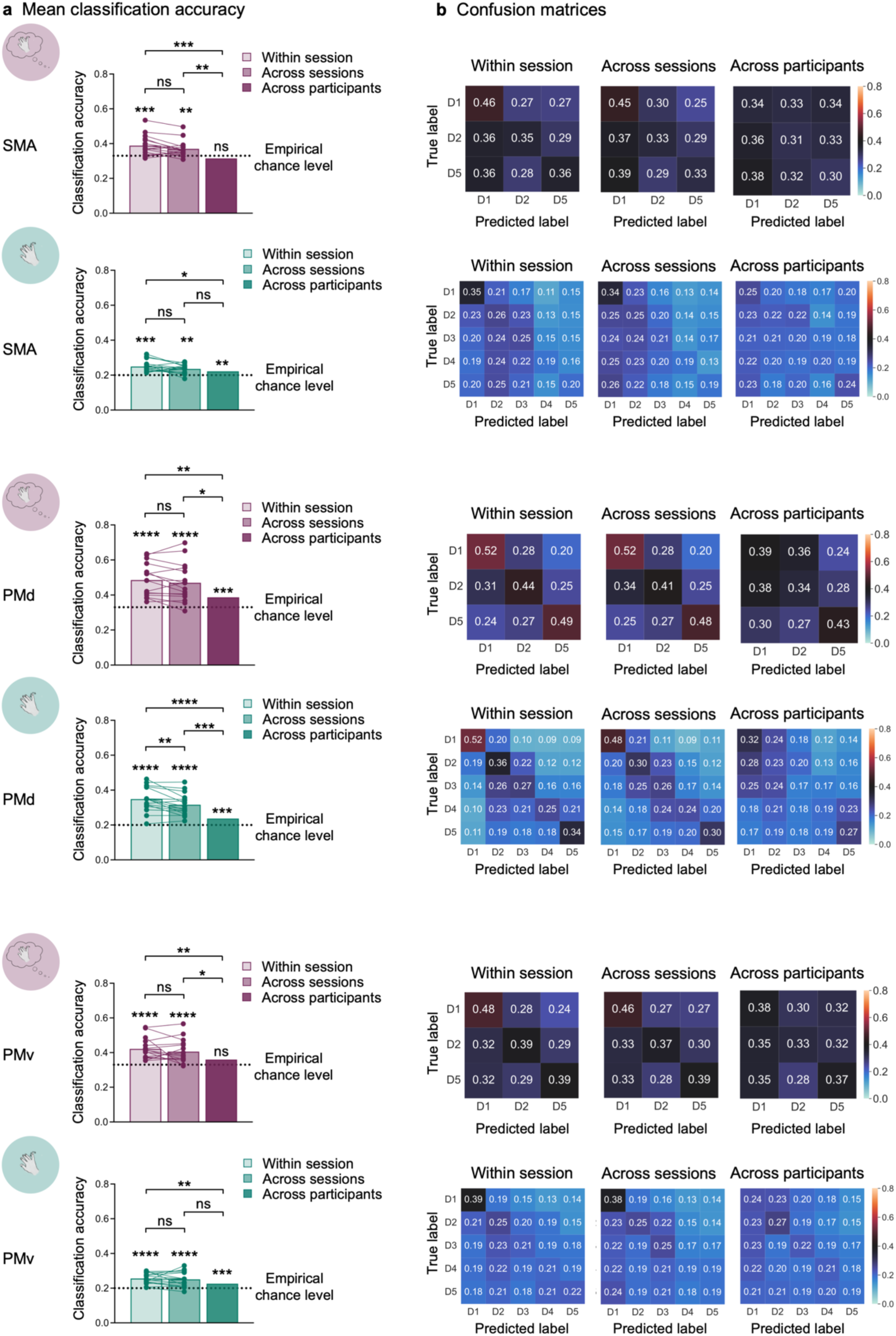
Decoding analysis of individual imagined and executed finger movements in secondary motor areas. **a**) Classification accuracy of five individual fingers during motor execution and three individual fingers during motor imagery in secondary motor areas. Within-session classification accuracy depicts the average accuracy of a leave-one-run-out cross-validation performed separately for each participant and session. Across-sessions is based on the average accuracy score of a within-participant classifier trained on all trials of one session and tested on all trials of the other session. Across-participants classification accuracy shows the average accuracy of a leave-one-participant-out cross-validation using the data of all participants. Asterisks on top of bars refer to the statistical difference of classification accuracy from the empirical chance level. **b**) Confusion matrices depicting the correctly classified trials (diagonal) and the misclassified trials (off-diagonal) in secondary motor areas (corresponding to the figures in the same row in **a**). Each row of the matrices refers to all trials of a finger, and the cells in a row show the % of predicted trials for each label. **** *p* < .0001; *** *p* < .001; ** *p* < .01; * *p* < .05; ns = non-significant. D1 (digit1) = thumb; D2 = index finger; D3 = middle finger; D4 = ring finger; D5 = little finger; SMA = supplementary motor area; PMd = dorsal premotor cortex; PMv = ventral premotor cortex.

**Supplementary Figure 7.**
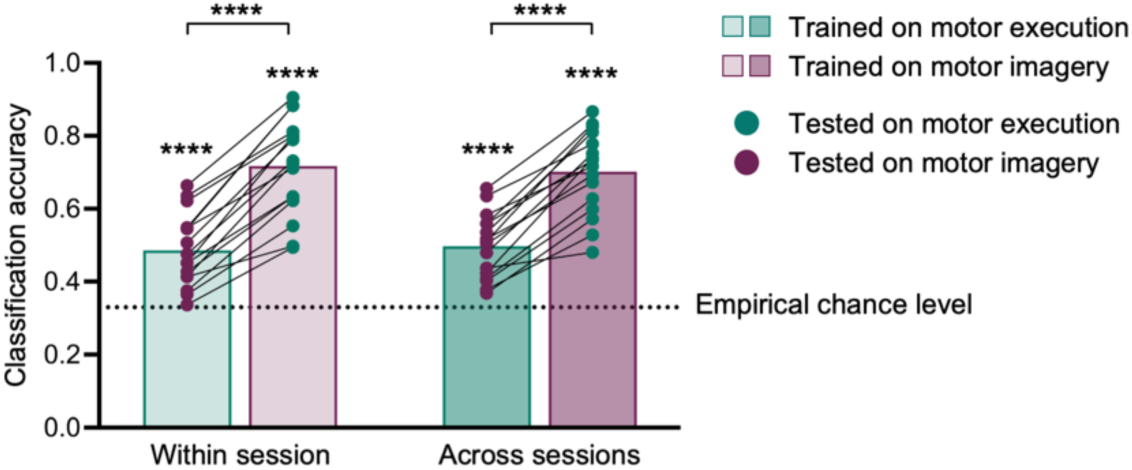
Across-task decoding analysis reveals neural similarity of finger representations activated through motor imagery and motor execution in the M1 hand area. Within-session classification accuracy depicts the average accuracy of a classifier trained on all data of one task (i.e., either motor imagery or motor execution) in one session and tested on all data of the other task in the same session, performed separately for each participant. Across-sessions shows the average accuracy of a classifier trained on all data of one task in one session and tested on all data of the other task in the other session. The significance bars refer to post-hoc contrasts based on the main effect of *Task* (motor imagery, motor execution: *F*_(1,45)_ = 321.62, *p* < .0001). There was no significant difference between within- and across-session classification accuracy (non-significant main effect *Comparison*: *F*_(1,45)_ = 0.03, *p* = .87) and no significant interaction effect (*F*_(1,45)_ = 1.20, *p* = .28). Asterisks on top of bars refer to the statistical difference of classification accuracy from the empirical chance level. The dotted lines represent the empirical chance level (i.e., 0.33) based on permutation testing. Dots depict data of individual participants. **** p < .0001; ns = non-significant.

**Supplementary Table 1a.** Verbatim instructions for fMRI sessions.

|  |
| --- |
| • Increase activity by **vividly imagining to move (feeling not seeing!) the instructed finger** of the right hand, for example by imagining:<br>• to move the finger up/down, left/right.<br>• to press a button, playing piano, typing on a keyboard.<br>• pulses in the muscle.<br>• ...<br>• Complex or forceful movements will give more activation than weak or simple movements.<br>• The imagined movement should **only involve the instructed finger** and no others. |
| • **Decrease activity for the not instructed fingers**, for example by imagining:<br>• that the other fingers were very cold, as if in a bucket of ice water.<br>• would not belong to the body or not exist at all.<br>• ... |
| • Imagine the movement as long as you can see the instruction on the screen.<br>• **Ensure to not actually move your fingers.** Also ensure no actual muscle activity anywhere in the body. Keep also your face muscles completely relaxed. |

**Supplementary Table 1b.** Self-reported strategies during session 1 and 2, separately for each participant. Please note that we only started to collect self-report of strategies in the fMRI session after study onset (from participant nr. 4 on).

| Participant | Session | Thumb | Index | Little |
| --- | --- | --- | --- | --- |
| 4 | 1 | pushing palm | pushing button | *did not recall* |
|  | 2 | pushing palm | brooding cheese | pushing button |
| 5 | 1 | "thumbs up" multiple time | holding up index finger | holding up little finger |
|  | 2 | typing | typing | typing |
| 6 | 1 | moving thumb up- and downwards,<br>pushing piano key,<br>typing,<br>moving thumb over surface | moving index up- and downwards,<br>pushing piano key,<br>typing,<br>moving index over surface | moving little finger up- and downwards,<br>pushing piano key,<br>typing,<br>moving little finger over surface |
|  | 2 | moving thumb up- and downwards,<br>pushing piano key,<br>typing,<br>moving thumb over surface | moving index up- and downwards,<br>pushing piano key,<br>typing,<br>moving index over surface | moving little finger up- and downwards,<br>pushing piano key,<br>typing,<br>moving little finger over surface |
| 7 | 1 | playing musical instrument,<br>scratching over table,<br>moving up and down in the air | playing musical instrument,<br>scratching over table,<br>moving up and down in the air | playing musical instrument,<br>scratching over table,<br>moving up and down in the air |
|  | 2 | moving it in a round circle, cutting the cake with one finger | moving it in a round circle, cutting the cake with one finger | moving it in a round circle, cutting the cake with one finger |
| 8 | 1 | moving finger to the left and right, spreading | typing | moving finger to the left and right, spreading |
|  | 2 | bending | typing | spreading |
| 9 | 1 | kneading bread, testing if it is ready to bake, making circles | looking through a map, pointing at landmark/location, making circles | making circles, imagining the circles and other fingers in ice |
|  | 2 | pressing my side, making circles, testing bread dough | looking through a map, pointing at landmark, making circles and lifting | lifting the little finger |
| 10 | 1 | counting number of guests with counting device, welcoming passengers | set timer on the oven | Groove of MRI |
|  | 2 | home button on phone, counting number of guests | set timer on the oven | typing on the edge of MRI |
| 11 | 1 | pressing a button | pressing a button | moving finger up and down |
|  | 2 | pressing a button | pressing a button | moving finger up and down<br>pressing a button (as alternative) |
| 12 | 1 | pressing on surface | pressing on surface | stretching outwards |
|  | 2 | pressing outwards | pressing on surface | stretching outwards |
| 13 | 1 | thumb game, twiddling the thumbs, à in sign language | ‘worm song’ with finger, making circles with index finger | ‘worm song’ with finger, making circles with little finger |
|  | 2 | à in sign language | ‘worm song’ | ‘worm song’ |
| 14 | 1 | thumb war, pressing controller button | pressing controller button, rotating a ring around using index finger, tapping on the desk | pinky swear, moving little finger around into a water bowl |
|  | 2 | extending my thumb thumb war, tapping on smartphone | sliding finger on surface, especially paper | pinky swear |
| 15 | 1 | rotating controller of a beamer, movements of pressing, extension, adduction, abduction and rotation | rotating controller of a beamer, movements of pressing, extension, adduction, abduction and rotation | movements of extension, adduction, abduction, rotation, pulling a hook downwards |
|  | 2 | rotating controller of a beamer, movements of pressing, extension, adduction, abduction and rotation, | rotating controller of a beamer, movements of pressing, extension, adduction, abduction and rotation, | movements of extension, adduction, abduction, rotation, pulling a hook downwards, pressing piano key |
|  |  | pressing spacebar on keyboard,<br>using gaming controller | pressing piano key and keyboard |  |
| 16 | 1 | making circles (like holding a joystick), up- and downwards movements, grabbing a handle | pulling a trigger, picking nose, scratching the head | moving finger left, right, up, down, pushing outwards, holding a rope |
|  | 2 | moving finger in figure of 8, up and down, apply sunscreen | Tickling, swiping a screen, moving finger in figure of 8 | moving finger to the side, swipe inside of a ham jar |

**Supplementary Table 2.** Group average of responses in visual analogue scales for session 1 and 2 and paired parametric or non-parametric tests.

| Variable | Mean (SD) Session 1 | Mean (SD) Session 2 | Test statistics | p-value (uncorr) |
| --- | --- | --- | --- | --- |
| Vividness | 66.75 (8.10) | 63.36 (10.30) | $t_{(15)} = 1.79$ | .09 |
| Motivation | 68.75 (14.79) | 64.69 (14.68) | $W = 111.00$ | .03 |
| Focus | 66.23 (14.68) | 59.07 (15.67) | $W = 106.00$ | .05 |
| Effort | 57.05 (12.89) | 59.95 (8.24) | $W = 68.00$ | 1.00 |

## Notes

### Competing Interest Statement

The authors have declared no competing interest.

